# Is there a need to implement standardisation into *in vitro* antimicrobial evaluation systems? A European collaboration perspective

**DOI:** 10.64898/2026.06.11.731574

**Authors:** Marie Attwood, Mark Brönstrup, Shampa Das, Hazel L. S. Fuchs, Pippa Griffin, Julien Lebrat, Benjamin Macklin, Sandrine Marchand, Derry K Mercer, Florian Michel, Alan Noel, Alina Nussbaumer-Proll, Markus Zeitlinger, Alasdair MacGowan

## Abstract

**Background:** Time kill curve (TKC) assessments are an essential step in the study of an antimicrobials pharmacodynamic characteristics. Surprisingly TKCs have not be formally standardised, therefore there remain concerns that different testing centres/methodologies may produce different results. Six centres participating in Gram-negative-Antibiotics NOW (GNA-NOW) consortium measured a series of TKCs with meropenem against *E. coli* to establish: Same-day (SD) vs different-day (DD) replication per centre (intra-site), and centre to centre (inter-site) correlations.

**Methods:** Meropenem was tested against three strains of *E. coli* (ATCC 25922; ESBL producer C1.55; OXA-48 producer C1.62). An inoculum of 1.5×10^6^ CFU was specified with meropenem concentrations of x0, x1 to x16 MIC; and sampling assessment of bacterial density was determined at 0-24h. Experiments were performed in triplicate, aerobically at 37°C. Centre-specific methodology was collected. Meropenem, media, bacterial strains, were shipped from one central laboratory to participating laboratories. ANOVA and Friedman tests were used to assess SD, DD and between centre replications.

**Results:** Assessment of the methodologies between centres revealed many differences, including bacterial inoculum, meropenem preparation, volume of TKC vessel, vessel materials, agitation vs static cultures and sampling volumes. Intra-centre SD and DD analysis for all strains were generally associated with *P*>0.05 suggesting consistency. Inter-centre SD and DD comparisons resulted in *P*<0.05, indicating variable total bacterial load measurement between centres.

**Conclusions:** TKC methodologies varied between different centres, and while intra-centre comparison of SD and DD were generally consistent, inter-centre comparisons were not. Standardisation of TKC methodologies is required.

## Introduction

The first successful administration of penicillin to treat bacterial infection was in 1941^1^which marked the beginning of the “golden era” for antibiotics. At the same time Harry Eagle was researching *in vivo* studies of penicillin activity, specifically the factors which impacted on antibiotic response, dose, and dosing interval^2 3^. Studies conducted by Eagle and others began the concept of anti-infective pharmacokinetic and pharmacodynamic (PK/PD) interactions.

In the 1980s, the continuation of *in vivo* studies produced experimental systems such as the neutropenic mouse-thigh infection model^4^ alongside the introduction of *in vitro* modelling systems, which allowed investigators to differentiate between the three classical PK/PD indices (C_max_, T>MIC and AUC/MIC). Since then, many similar studies have identified relevant PK/PD indices across a range of antimicrobial classes^5^. PK/PD parameters have been used for antimicrobial development and pre-clinical/ clinical studies of antimicrobials^6^. With the rise of antimicrobial resistance (AMR), there has been an increased number of initiatives launched to tackle this issue. One such initiatives is the GNA NOW project which is part of the IMI AMR Accelerator programme^7^.

The GNA NOW project aims to address the urgent need for antibiotics to treat Gram-negative infections. The consortium is comprised of 11 independent partners consisting of industry, small and medium sized enterprises, universities, research organisations, public bodies, and non-profit groups. While the consortium benefits from a diverse range of skills and expertise, there is a general overlap of practical microbiology techniques i.e. time-kill curve (TKC) studies, *<u>in vitro</u>* modelling simulations/systems and *<u>in vivo</u>* models.

*In vitro* Time Kill Curves (TKCs), *in vitro* modelling simulations (IVMs), and *in vivo* murine infection model data are frequently used to support suggested dosing strategies in submissions to regulatory authrorites^8,9^. To date, Time kill curves (TKCs) are considered the gold standard of *in vitro* pharmacodynamic (PD) evaluation of anti-infectives versus bacterial growth, allowing the pattern of kill for an antibacterial to be established^10^. However, methodologies vary in several areas, such as materials, bacterial growth phases, drug preparation, culture vessel volume, and inoculum.

These variations prompted us to explore the reproducibility of *in vitro* TKCs, focusing on two main questions: Are same-day (SD) TKC replications comparable with different-day (DD) replications within individual centres (intra-site)? And secondly, do we get result consistency across multiple centres (inter-site) where various methodologies are employed?

## Materials and Methods

Supplementary Figure 1 illustrates the evaluation workflow and progression markers which was employed for this study.

### Bacterial isolates

Three *E. coli* strains were tested, including the CLSI & EUCAST reference strain *E. coli* ATCC 25922 (meropenem MIC = 0.03 mg/L) and two clinical isolates with known resistance mechanisms. Strain C1.62 is an ESBL producer (meropenem MIC = 0.03 mg/L) and C1.56 is an OXA-48 producer (meropenem MIC = 1 mg/L). Antibiograms can be found in supplementary Table S1.

The target inoculum was set at 1.5 × 10^6^ CFU/mL to mirror bacterial inocula commonly used in *<u>in vivo</u>* pharmacodynamic models.

### Antimicrobial

Meropenem was selected for this study as it has been studied extensively *in vitro* and is often used as the gold standard treatment for hard-to-treat bacterial infections. We sourced a pharmaceutical preparation from ACS Dobfar: 1 g powder for solution for injection/infusion. Solvent and diluent stated by manufacturer: H_2_O. Potency 100%. Targets were set as multiples of MIC; x0 (growth control), x1, x2, x4, x8, x16.

### Media

Growth medium employed was Muller Hinton Broth II – BD BBL™ Dehydrated culture Media, also known as cation-adjusted Mueller Hinton broth (CA-MHB). Certificates of compliance were BS12, CLSI, CMPH2, MCM9 and the manufacturers reconstitution method using deionised H_2_O was carried out, followed by sterilisation via autoclaving. Environmental conditions stated for TKC incubations were aerobic conditions at 37^°^C.

### Experimental data points tested

Three replicates per site were performed on the same day, and then subsequent replicates on 3 different days. Timepoints for bacterial count enumeration were at hours 0, 2, 4, 6, 8 and 24 post-inoculation. All results were reported in CFU/mL.

### Experimental sites

Listed in alphabetical order; Bioaster Fondation De Cooperation Scientifique, Lyon, France. Helmholtz Centre for Infection Research, Braunschweig, Germany. Medizinische Universitaet Wien, Vienna, Austria. North Bristol NHS trust, Bristol, United Kingdom.

Universite De Poitiers, Poitiers, France. University of Liverpool, Liverpool, United Kingdom.

## Results

### Site methodology variations

There were major differences in methodology across sites, as described in Table 1. TKC culture vessel, the total volume of media in that vessel, inoculum state (i.e. prepared from exponentially growing or stationary phase cultures), inoculum added, meropenem preparation, sample volume for CFU enumeration and CFU enumeration itself all varied across the centres. The following TKC experiments were designed to evaluate whether differences in methodologies (Table 1) led to inconsistent data points, reveal systematic errors, and introduce bias that could affect data interpretation.

**Table 1.** Variations in experimental procedures across sites.

| Description of action | Site A | Site B | Site C | Site D | Site E | Site F |
| --- | --- | --- | --- | --- | --- | --- |
| Method | Culture vessels | Microbroth | Microbroth | Culture vessels | Culture vessels | 96 well plate |
| Inoculum - Target 1.5 x 10E6 |  |  |  |  |  |  |
| Preparation | Overnight culture | Overnight culture | Overnight culture | Overnight culture | Overnight culture | - |
| Volume of culture | 10 mL | - | 10 mL | 10 mL | 20 mL | - |
| Culture conditions | 37°C, aerobic, Static | - | 37°C, aerobic, under stirring conditions | 37°C, aerobic, static | 37°C aerobic under stirring conditions | 37°C aerobic under stirring conditions |
| Diluent for culture | MHB II broth | Sodium chloride 0.9% | MHB II broth | MHB II broth | MHBII broth | - |
| Dilution | 1:9 | 1:100 | 01:50 | 1:9 | - | 01:10 |
| Bacterial phase | Lag | Lag | - | Lag | Log / Lag | Log |
| Volume of inoculum added to test vessel | 1000 µL | 30 µL | 100 µL | 1000 µL | - | 3-300 µL |
| Media - MHBII (BD) |  |  |  |  |  |  |
| Volume | 9000 µL | 3040 µL | 1000 µL | 9000 µL | - | 29000 µL |
| Other additional information | Media prewarmed 24 hours before test start | - | - | Prewarmed 24 hours before test start | - | - |

| Description of | Site A | Site B | Site C | Site D | Site E | Site F |
| --- | --- | --- | --- | --- | --- | --- |

| action |  |  |  |  |  |  |
| --- | --- | --- | --- | --- | --- | --- |
| **Drug - Meropenem** |  |  |  |  |  |  |
| **Preparation** | Aqueous | Aqueous | Aqueous | Aqueous | PBS | Aqueous |
| **Diluent** | MHB II broth | H<sub>2</sub>O | MHB II broth | MHB II broth | - | MHB II broth |
| **Dilution series method** | Individual for each culture vessel | - | Serial dilution | Individual for each culture vessel | - | Individual for each culture vessel |
| **Stock volume for test method** | Stock levels are generated on specific drug concentration required for each Individual culture vessel | 30 µL | 1000 µL (added to MHBII 1000 µL, serial dilution 1000 µL, total volume = 1000 µL) | Stock levels are generated on specific drug concentration required for each Individual culture vessel | - | 1:10 - 1:100 |
| **Assessment of drug concentrations** | - | Not stated | Not stated | Ascertained via HPLC on site | - | - |
| **Storage of samples for drug concentration assessment** | - | Not stated | Not stated | At -80°C | - | -80°C |
| **Additional information** |  |  |  |  |  |  |
| **TKC conditions** | Culture vessels incubated at 37°C±1, aerobically and static | Not stated | Microbroth plate incubated at 37°C under agitation | Culture vessels incubated at 37°C±1, aerobically and static | - | Microbroth plate incubated at 37°C under agitation |
| **Total volume of TKC** | 10 mL | 400 µL | 200 µL | 10 mL | 30 mL | 100 µL |
| **Description of action** | Site A | Site B | Site C | Site D | Site E | Site F |
| CFU determination |  |  |  |  |  |  |
| **Method** | Eddie Jet | Via agar plate | Via agar plate | Vial Don Whitley Spiral platter (WASP) | - | Via agar plate |
| **Additional information** | Neat samples (50µL) spiralled onto agar plates using Log function. 10<sup>3</sup> dilutions performed (saline diluent) to give a range up to 4.0x10<sup>8</sup> CFU) |  | Using 96-well plate with 0.9% NaCl. 1/10 serial dilution, assess CFU visually. Plate 1/3 on Agar plate | Neat samples (50µL) spiralled onto agar plates using Log function. 10<sup>3</sup> dilutions performed (saline diluent) to give a range up to 4.0x10<sup>8</sup> CFU) | - | OD450 value, CFU count, dilution factor and CFU/mL for each time point |
| **Sampling Volume** |  | 20 µL | 100 µL | 100 µL | - | Unknown |
| **Time points** | 0,2, 4, 6, 24 | 0, 2, 4, 6, 8, 24 | 0, 2, 4, 6, 8, 24 | 0, 2, 4, 6, 8, 24 | - | 0, 2, 4, 6, 8, 24 |
| **Incubation time** | 16-20 hours | Overnight | 16-24 hours | 16-20 hours | - | 24 hours |
| **Atmosphere** | Incubated at 37°C±2 | 37°C, aerobic | Incubated at 35°C | Incubated at 37°C±1 | - | Incubated at 37°C±1 |
| **EoR** | - | - | - | - | - | - |
| **Isolates saved** | - | Not stated | Not stated | Yes | - | - |
| **Method of storage** | - | Not stated | Not stated | Microbank cryovial beads | - | - |

Figure 1 shows the TKC Meropenem versus ATCC 25922 *E. coli* CFU trends observed for same day replications for all six centres. x4 to x16 MIC multiple bioburdens exhibit similar trends across all sites. However, there is visible variation in the trends observed for the growth control x1 and x2 MIC multiples over the 24-hour period. The bioburden trends for *E. coli* bacterial strains C1.55 and C1.62 are similar. This suggests that the observed variations in bacterial kill and regrowth are not bacteria-drug specific but may instead stem from methodological differences in data collection.

**Figure 1.**
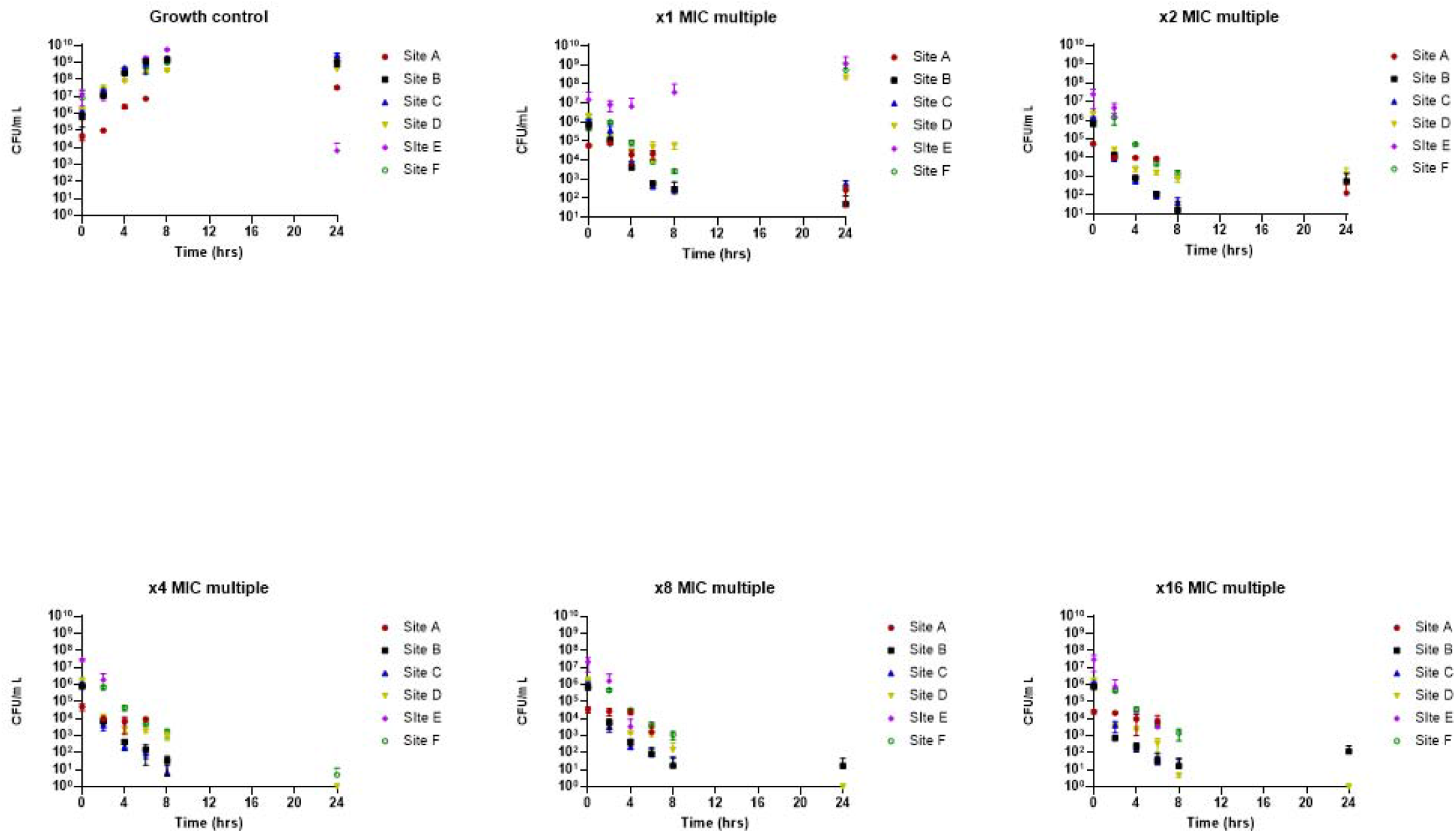
Sites A -F, ATCC *E. coli* 25922 same day comparison.

Figure 2 shows ATCC 25922 *E. coli* CFU trends observed for different day replications per site for timepoints 0, 4, and 24. Here you can see that for each MIC multiple replications within each site, the data points are generally consistent. However, variations across different sites are also observable. This also supports that theory that variations in methodology are contributing to systemic bias. All CFU data for all 3 strains can be found in supplementary Table S2 and S3.

**Figure 2.**
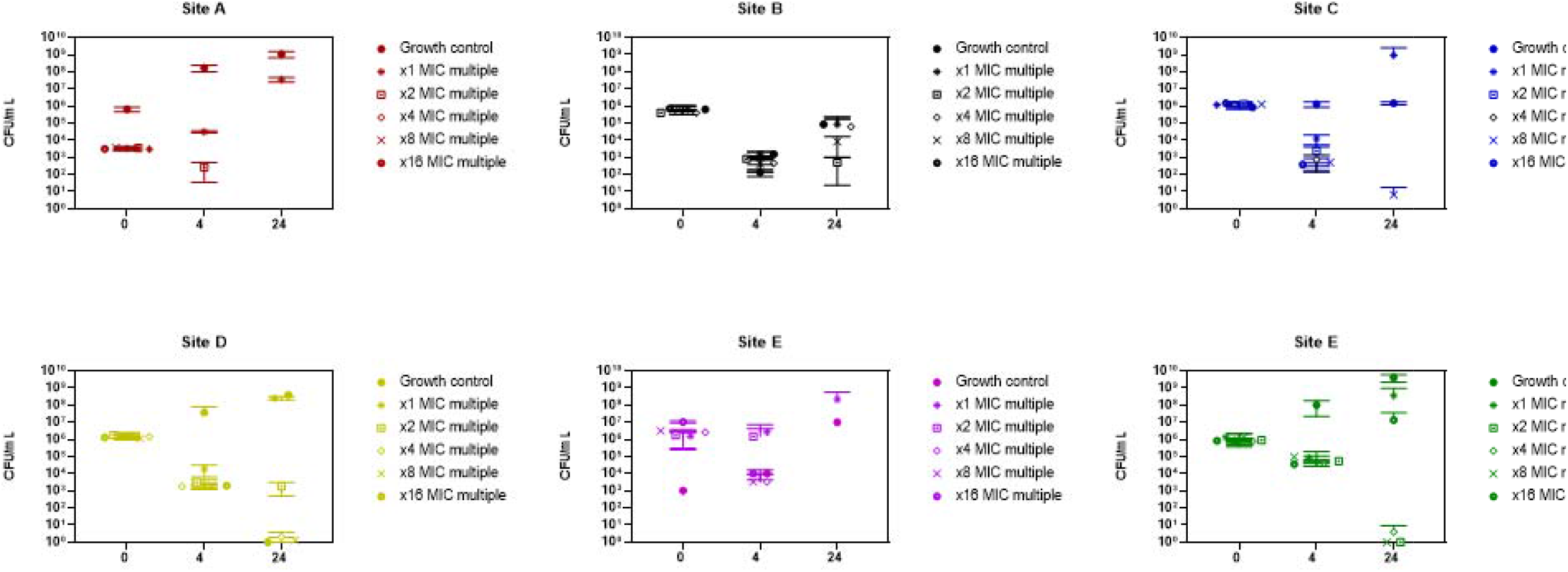
Sites A – F, ATCC *E. coli* 25922 different day comparison.

### Description of the data

Generally, the kill/growth trends across all centres for same day replications demonstrate visually similar patterns of growth or kill despite a range of initial inoculums. Meropenem concentration has an inverse relationship with CFU/mL, however there are some discrepancies at x1 MIC multiple meropenem concentration at timepoint 24 hours (Figure 1). The different day data (Figure 2) shows on visual inspection increased spread of data points particularly for GC and x1 multiplies but overall similar kill/growth trends. The data from the other two bacterial strains were also plotted and showed similar kill/growth trends and are shown on Supplementary Tables S4 and S5, suggesting that this is a not a bacterial strain phenomenon but rather a methodological one.

### Data analysis

#### Statistical tests and assumptions used

To compare the distribution between inoculum repeats, an analysis of variance two-way ANOVA statistical test was used. The assumptions are that the data is normally distributed for same day versus different day intra-site replications and also inter-site replications. Target bacterial inoculum at timepoint 0 (T0) was set at 1.5×10^6^ CFU/mL to establish baseline CFU/mL prior to antibiotic exposure. The 1^st^ Null hypothesis, H_0_, is the expectation that repeated measures at T0 for intra-site data is consistent. The alternative hypothesis, H_1_, is that the repeated measures at T0 intra-site (across MIC multiplies) are statistically different. The second null hypothesis, H_0_, is that repeated measures at T0 inter-site is consistent. The alternative second null hypothesis, H1, is that repeated measures at T0 inter-site is statistically different.

Normal distribution cannot be assumed once bacteria have been exposed to an antibiotic, such as meropenem, and therefore another statistical test was used for these evaluations.

Friedman test was employed to determine any statistical difference between intra and inter centre same day and different day data. In this application of the Friedman test, the null hypothesis, H_0_, is the expectation that concentration of antibiotic has no effect on the trend of total number CFU/mL. Therefore, the alternate hypothesis, H_1_, states that antibiotic concentration affects the trend of CFU/mL. To assess the rate of bacterial kill, the analysis focused on T4 and T24 time points. Friedman test values at timepoints previously specified were compared against the threshold value derived from Chi-squared (χ^2^) distribution tables. If the Friedman test values exceed this threshold, it is indicative that the replications within individual sites and across sites are consistent with each other. To summate this the higher the test statistic which exceeds the threshold the more consistent the data points. The number of experiments using different antibiotic concentrations in each group, k, was 6, so the reported χ^2^ (DoF = 5, a = 0.05) threshold value was Fr_threshold_ = 11.070.

Table 2 describes same day analysis per centre, different day analysis per centre and same day and different day between centre analysis. For most of the centres there are no significant differences (green highlight) for same day or different day replications within a centre for the three *E. coli* strains. Hence, inoculum variation is determined by centre and not influenced by bacterial strain. In contrast, for all the centres the cross-centre replications on the same day and on different days were significantly different, indicating that the bacterial inocula were not consistent across the centres. (Table 2). This suggests that individual centre methodological inoculum practices, (such as sample size) has a significant impact on achieving target CFU and/or CFU enumeration reproducibility rather than strain-dependent differences.

**Table 2.** Summary analysis for same day and different day replications for both intra and inter-centre initial inoculum for three *E. coli* strains at T0 using two-way ANOVA.

| Site A |  |  |  |  |  |  |
| --- | --- | --- | --- | --- | --- | --- |
| ANOVA Table | 25922 |  | C1.55 |  | C1.62 |  |
|  | P value | Significant | P value | Significant | P value | Significant |
| Per site SD | 0.0707 | No | - | - | 0.0794 | No |
| Per site DD | 0.002 | Yes | - | - | 0.0004 | Yes |
| Cross site SD | <0.0001 | Yes | - | - | <0.0001 | Yes |
| Cross site DD | <0.0001 | Yes | - | - | <0.0001 | Yes |
| Site B |  |  |  |  |  |  |
| ANOVA Table | 25922 |  | C1.55 |  | C1.62 |  |
|  | P value | Significant | P value | Significant | P value | Significant |
| Per site SD | 0.9867 | No | 0.1204 | No | 0.2041 | No |
| Per site DD | 0.8031 | No | 0.6461 | No | 0.1191 | No |
| Cross site SD | <0.0001 | Yes | <0.0001 | Yes | <0.0001 | Yes |
| Cross site DD | <0.0001 | Yes | <0.0001 | Yes | <0.0001 | Yes |
| Site C |  |  |  |  |  |  |
| ANOVA Table | 25922 |  | C1.55 |  | C1.62 |  |
|  | P value | Significant | P value | Significant | P value | Significant |
| Per site SD | 0.0523 | No | >0.9999 | No | 0.6234 | No |
| Per site DD | 0.2625 | No | 0.3241 | No | 0.5562 | No |
| Cross site SD | <0.0001 | Yes | <0.0001 | Yes | <0.0001 | Yes |
| Cross site DD | <0.0001 | Yes | <0.0001 | Yes | <0.0001 | Yes |
| Site D |  |  |  |  |  |  |
| ANOVA Table | 25922 |  | C1.55 |  | C1.62 |  |
|  | P value | Significant | P value | Significant | P value | Significant |
| Per site SD | 0.5787 | No | 0.7715 | No | 0.9625 | No |
| Per site DD | 0.9863 | No | 0.3312 | No | 0.5311 | No |
| Cross site SD | <0.0001 | Yes | <0.0001 | Yes | <0.0001 | Yes |
| Cross site DD | <0.0001 | Yes | <0.0001 | Yes | <0.0001 | Yes |
| Site E |  |  |  |  |  |  |
| ANOVA Table | 25922 |  | C1.55 |  | C1.62 |  |
|  | P value | Significant | P value | Significant | P value | Significant |
| Per site SD | <0.0001 | Yes | <0.0001 | Yes | 0.8892 | No |
| Per site DD | <0.0001 | Yes | <0.0001 | Yes | <0.0001 | Yes |
| Cross site SD | <0.0001 | Yes | <0.0001 | Yes | <0.0001 | Yes |
| Cross site DD | 0.0052 | Yes | <0.0001 | Yes | <0.0001 | Yes |
| Site F |  |  |  |  |  |  |
| ANOVA Table | 25922 |  | C1.55 |  | C1.62 |  |

|  | P value | Significant | P value | Significant | P value | Significant |
| --- | --- | --- | --- | --- | --- | --- |
| Per site SD | 0.6943 | No | 0.3041 | No | 0.4202 | No |
| Per site DD | 0.4311 | No | 0.004 | Yes | 0.0042 | Yes |
| Cross site SD | <0.0001 | Yes | <0.0001 | Yes | <0.0001 | Yes |
| Cross site DD | <0.0001 | Yes | <0.0001 | Yes | <0.0001 | Yes |

### T2+T4 and T24 same day analysis

Two hour and four-hour timepoints were used to assess kill, and eight hour and twenty four-hour timepoints were used to determine any regrowth trends. All Friedman test statistics were compared to a threshold value of 11.07 χ^2^, and are shown on Figure 3 and supporting supplementary Tables S4 and S5.

**Figure 3.**
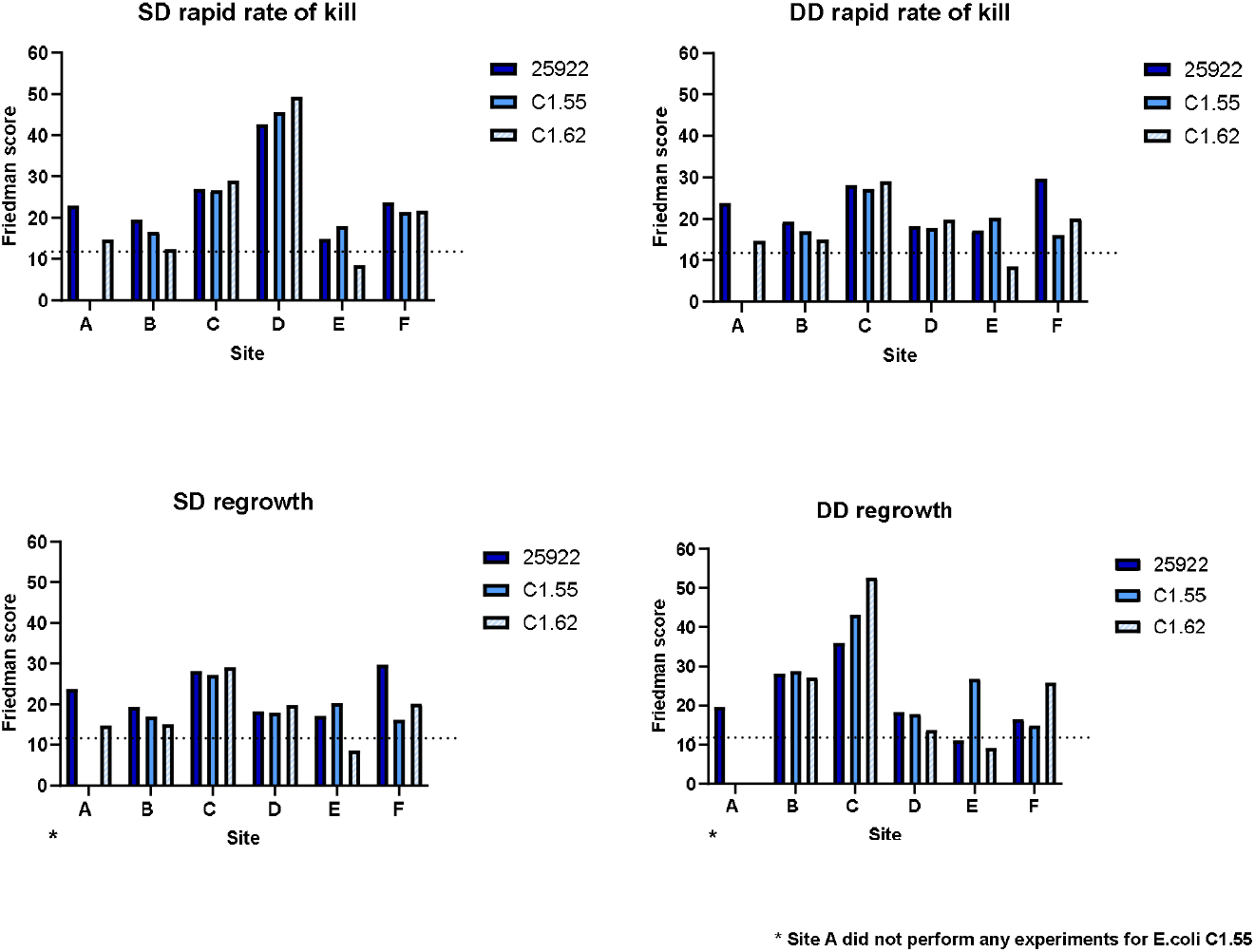
Intra -centre same day (SD) and different day (DD) comparisons (dotted line represents the threshold value)

Intra-centre comparison shows that generally bacterial kill and regrowth patterns are consistent per centre for same day or different day replications and also for the three bacterial strains, as most Friedman scores exceed the threshold value. Inter-centre comparison of same day, different day, rapid kill and regrowth assessment can be seen in Figure 4.

**Figure 4.**
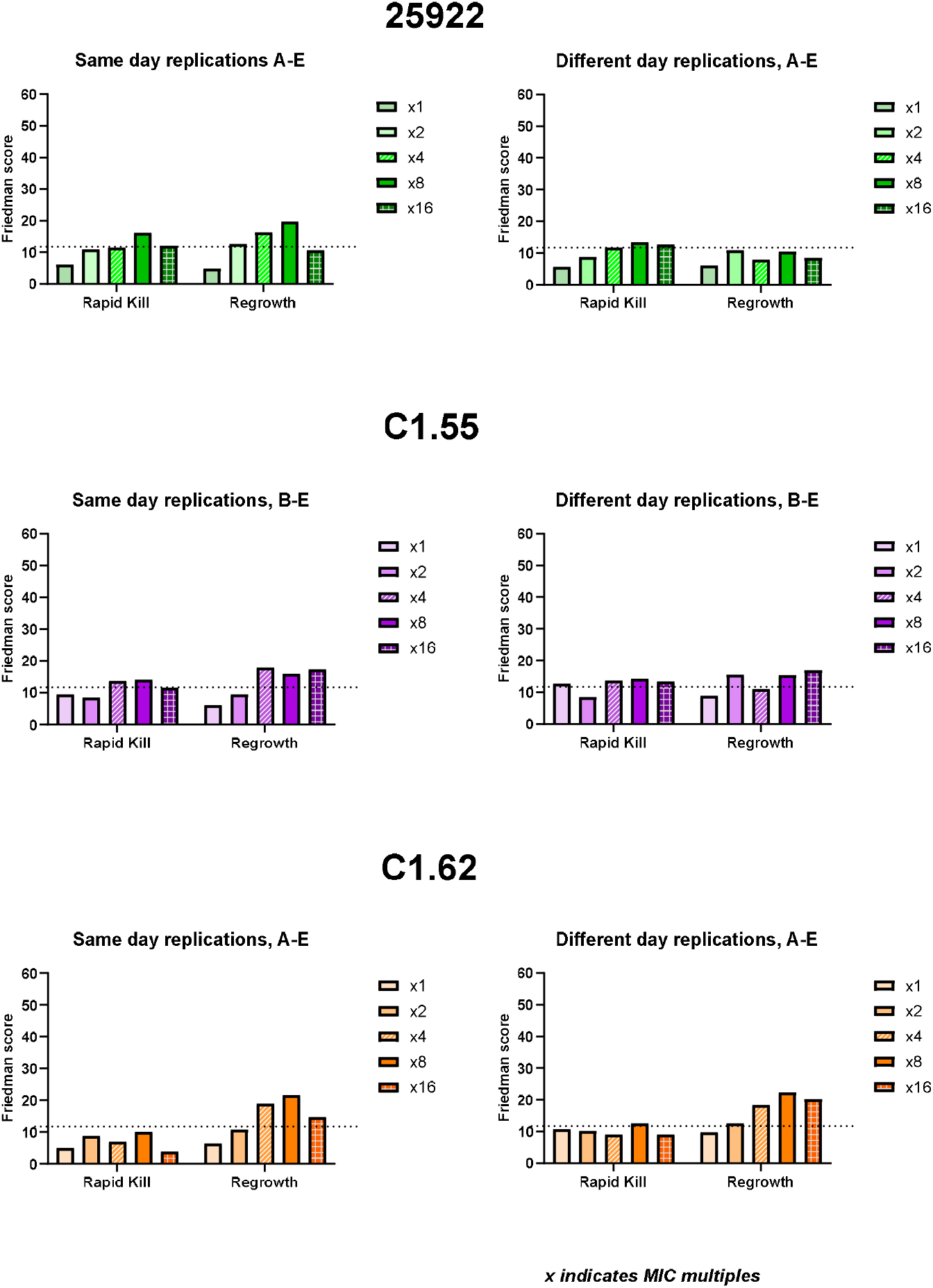
Inter-site same day and different day comparisons for *E. coli* ATCC 25922, C1.55 and C1.62 (dotted line represents the threshold value)

Inter-centre comparisons demonstrate that bacterial kill and growth patterns are inconsistent across sites irrespective of same day or different day replications and bacterial strain. This is shown by most Friedman scores falling below the threshold value.

Figure 5 shows data analysis from sites C and D only. These sites were selected as total bacterial counts at T0 which were statistically consistent between for SS, DD, intra and inter-site comparisons. Despite reproducible inoculum there are still statical differences CFU trends for initial rate of kill and regrowth.

**Figure 5.**
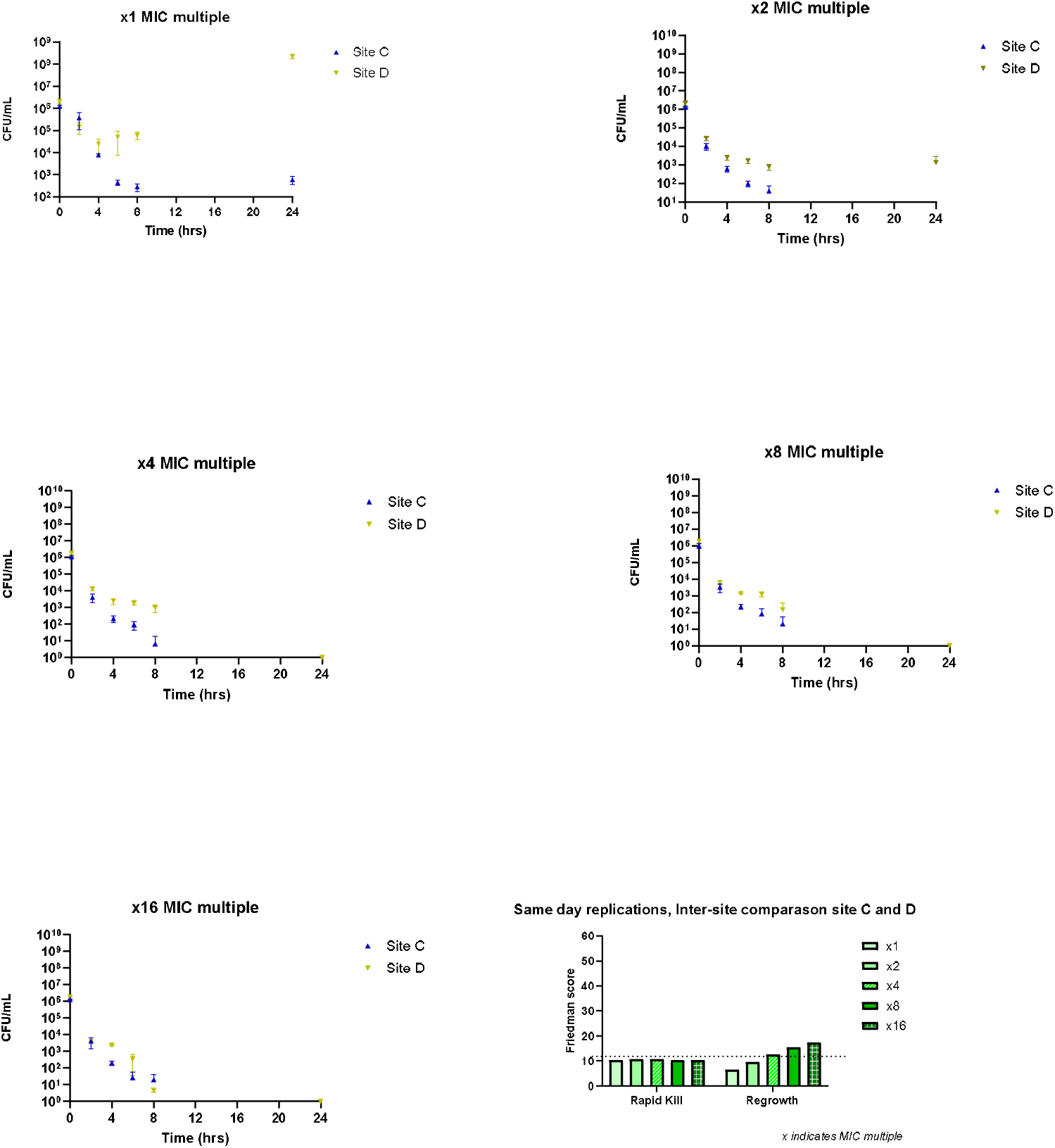
Summary graphs *E. coli* ATCC 25922 CFU/mL and inter-site Friedman scores for sites C and D only.

## Discussion

Previous comparative publications have demonstrated that initial inoculum plays a key factor in analysis of changes in bacterial load (Dalhoff et al 2020^11^) in *in vitro* modelling systems.

This study compared eight types of systems (closed and open) and established that CFU washout was not a factor in population analysis profiles, but rather that the starting inoculum affected results significantly. Unsurprisingly, this also appears to be true for this study, as we have observed disparity for some intra-site and inter-site replications. Inocula variation, appears to not be bacterial strain dependant and is likely attributed from differences in; the size of the culture vessel, volume of inoculum used, sample volume taken for CFU enumeration, or a combination of all these factors. Centres C and D demonstrate consistent inoculum target obtainment for both same day and different day replications. However, there was still statistical variation in rapid kill and regrowth. This implies that additional factors may be contributing to the observed variations in bacterial growth trends.

We have established that rapid kill and regrowth trends for intra-site same day replication was statistically consistent, and this generally was also the case for intra-site different day replications. This indicates that the TKC experiments are reproducible per institution.

However, for both same day replications and different day replications, we see statistical differences in kill or regrowth trends between the centres. The cause of variation could be attributed to methodological differences. The methodology used for the TKC varied in almost every aspect across the six centres, but can categorised into two groups, i.e. volume of culture and culture conditions. The volume of TKC culture itself varied from 100 µL to 30 mL, and sampling volume size from the TKC culture varied from 20 µL to 100 µL. There was also variation in culture conditions, specifically between use of static or agitated cultures. There are publications where agitation has led to promotion of biofilm formation^12^, which alters PK/PD interactions.

## Conclusion

The findings of the study have implications for multisite research collaborations established to support antimicrobial drug development. Even for the well-established and seemingly ‘routine’ TKC experiment, experimental protocols need to be defined in detail including many technical factors from the outset. If this is too complex a process, then a central laboratory for all TKC experiments seems more advisable than decentralised testing. Moreover, often TKCs are used to inform experimental design of in the in vitro modelling systems, which are also not standardised. Existing CLSI^8^ and ECRAID^13^ guidelines do not specifically address these technical details, likely to maintain flexibility in experimental concepts. However, there is an increasing demand for standardization in all aspects of research^14^ ensures efficiency, enhances credibility, facilitates collaboration which all supports meta-analysis enabling pooling of data from multiple sources deriving stronger conclusions. We recognize that standardisation poses challenges, with the most significant concern being its potential to hinder innovation. Therefore, a balanced approach, opposed to a rigid process should be adopted to allow for thoughtful adaptation.

Consequently, we propose that the following elements which should be considered and standardised upon project commencement: Inoculum targets set with limits of confidence, TKC culture size and agreement on environmental conditions. These factors should be specific to the test compound and organism of interest and provide project harmonisation.

## Acknowledgement

The GNA NOW consortium has received funding from the Innovative Medicines Initiative 2 Joint Undertaking under Grant Agreement n°853979. This Joint Undertaking receives the support from the European Union’s Horizon 2020 research and innovation programme and EFPIA.

## Disclaimer

Funded by the European Union, the private members, and those contributing partners of the IMI JU. Views and opinions expressed are however those of the author(s) only and do not necessarily reflect those of the aforementioned parties. Neither of the aforementioned parties can be held responsible for them.

## Transparency Declarations

MA, PG, ARN and APM holds research grants/activities with Merck, Shionogi, InfectoPharm, GSK, Roche, BioVersys, VenatoRx Pharmaceuticals, iFAST, Technomede, Oxford Drug Design, JPIAMR and NIHR; APM provides consultancy advice to Shionogi, Roche, Bicycle Therapeutics and BioVersys. DM provides consultancy advice to INCATE, Selmod GmbH and University of Basilicata. All other authors declare no competing interests

