## Supplementary material for "Is there a need to implement standardisation into *in vitro* antimicrobial evaluation systems? A European collaboration perspective": TKC Quality manuscript 1of 2_supplimentary .docx

**Supplementary information**

**Figure S1 - Evaluation workflow for cross site time kill curve reproducibility evaluation.**

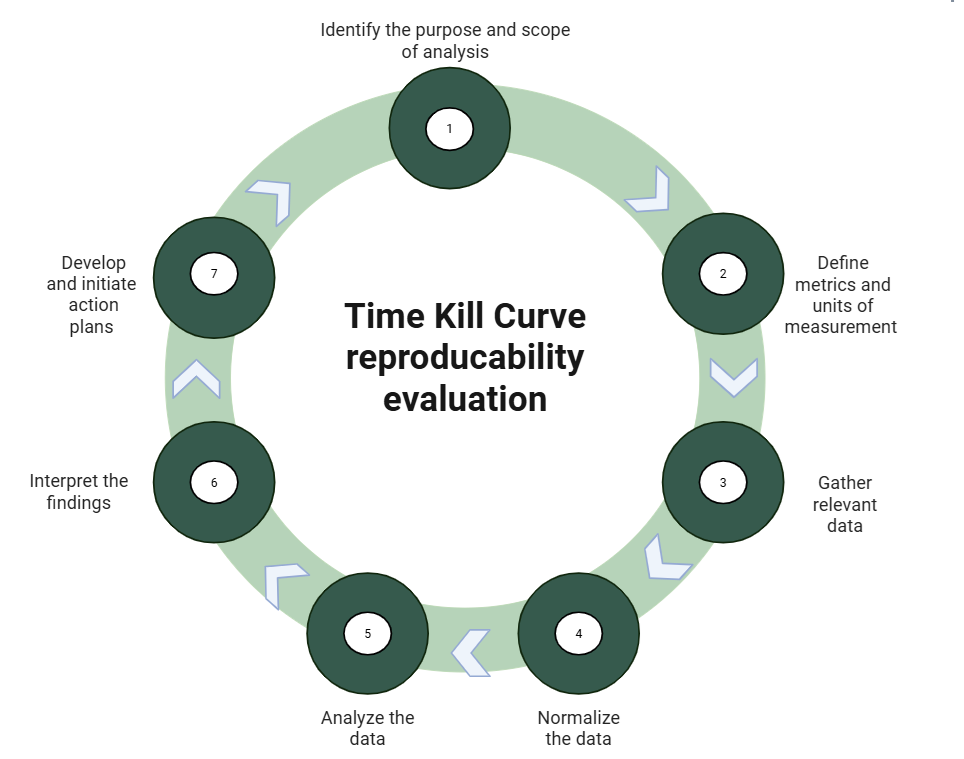

Table S1 – Antibiogram of *E. coli* strains

| Antimicrobial | mg/L | | |
| --- | --- | --- | --- |
|  | 25922 | C1.55 | C1.62 |
|  | ATCC | ESBL | OXA-48 |
|  | *E. coli* | *E. coli* | *E. coli* |
| Amikacin | 1 | 4 | 2 |
| Ceftazidime Avibactam | 0.12 | >16 | 8 |
| Ciprofloxacin | 0.008 | >1 | >1 |
| Colistin | 0.25 | 0.5 | 1 |
| Fosfomycin | 0.5 | 0.25 | 0.25 |
| Meropenem | 0.015 | 0.03 | 1 |
| Tigecycline | 0.03 | 0.25 | 0.25 |

Table S2 – Same day source data all strains

| Same day replications | | | | | | | | | | | | | | | | | | |
| --- | --- | --- | --- | --- | --- | --- | --- | --- | --- | --- | --- | --- | --- | --- | --- | --- | --- | --- |
| *E. coli* 25922 Site A | | | | | | | | | | | | | | | | | | |
| Timepoint | GC | | | x1 | | | x2 | | | x4 | | | x8 | | | x16 | | |
|  | Mean | SD | N | Mean | SD | N | Mean | SD | N | Mean | SD | N | Mean | SD | N | Mean | SD | N |
| 0 | 44000 | 18330.3 | 3 | 58666.67 | 8326.66 | 3 | 56000 | 8000 | 3 | 49333.33 | 20132.9 | 3 | 34666.67 | 12858.2 | 3 | 25333.33 | 6110.1 | 3 |
| 2 | 101333.3 | 8326.66 | 3 | 77333.33 | 10066.4 | 3 | 10400 | 800 | 3 | 10000 | 2116.6 | 3 | 26666.67 | 12220.2 | 3 | 21333.33 | 2309.4 | 3 |
| 4 | 2533333 | 611010 | 3 | 20000 | 14422.2 | 3 | 10000 | 1385.64 | 3 | 6666.667 | 5460.16 | 3 | 24000 | 6928.2 | 3 | 9333.333 | 8326.66 | 3 |
| 6 | 7600000 | 400000 | 3 | 21333.33 | 10066.4 | 3 | 9066.6667 | 1006.64 | 3 | 9333.333 | 832.666 | 3 | 1600 | 2116.6 | 3 | 6666.667 | 8326.66 | 3 |
| 8 | - | - | - | - | - | - | - | - | - | - | - | - | - | - | - | - | - | - |
| 24 | 34666667 | 6110101 | 3 | 266.6667 | 230.94 | 3 | 133.33333 | 230.94 | 3 | 0 | 0 | 3 | 0 | 0 | 3 | 0 | 0 | 3 |
| *E. coli* 25922 Site B | | | | | | | | | | | | | | | | | | |
| Timepoint | GC |  |  | x1 |  |  | x2 |  |  | x4 |  |  | x8 |  |  | X16 |  |  |
|  | Mean | SD | N | Mean | SD | N | Mean | SD | N | Mean | SD | N | Mean | SD | N | Mean | SD | N |
| 0 | 630000 | 467654 | 3 | 666667 | 256580 | 3 | 666667 | 175594 | 3 | 750000 | 50000 | 3 | 683333 | 28868 | 3 | 733333 | 28868 | 3 |
| 2 | 11833333 | 2753785 | 3 | 116667 | 20817 | 3 | 13833 | 4368 | 3 | 6667 | 1155 | 3 | 5667 | 3253 | 3 | 717 | 126 | 3 |
| 4 | 2.45E+08 | 3.9E+07 | 3 | 4167 | 577 | 3 | 833 | 126 | 3 | 417 | 104 | 3 | 417 | 152.753 | 3 | 233 | 126 | 3 |
| 6 | 1.2E+09 | 1.8E+08 | 3 | 600 | 132 | 3 | 117 | 29 | 3 | 150 | 132 | 3 | 83 | 104.079 | 3 | 33 | 58 | 3 |
| 8 | 1.5E+09 | 5E+07 | 3 | 300 | 435.886 | 3 | 16.673333 | 28.8617 | 3 | 33 | 28.8617 | 3 | 17 | 28.8617 | 3 | 16.67333 | 28.8617 | 3 |
| 24 | 1E+09 | 4.3E+08 | 3 | 50.00667 | 87 | 3 | 566.67 | 814.449 | 3 | 0 | 0 | 3 | 16.67333 | 28.8617 | 3 | 117 | 125.826 | 3 |
| *E. coli* 25922 Site C | | | | | | | | | | | | | | | | | | |
| Timepoint | GC | | | x1 | | | x2 | | | x4 | | | x8 | | | x16 | | |
|  | Mean | SD | N | Mean | SD | N | Mean | SD | N | Mean | SD | N | Mean | SD | N | Mean | SD | N |
| 0 | 1206667 | 231805 | 3 | 1206667 | 160416 | 3 | 1286666.7 | 113725 | 3 | 1113333 | 144684 | 3 | 1020000 | 321870 | 3 | 1286667 | 61101 | 3 |
| 2 | 23333333 | 1331666 | 3 | 369333.3 | 261276 | 3 | 10333.333 | 3910.67 | 3 | 4160 | 2157.04 | 3 | 3380 | 1777.98 | 3 | 4100 | 2666.46 | 3 |
| 4 | 4.77E+08 | 2.9E+07 | 3 | 7666.667 | 1026.32 | 3 | 626.66667 | 213.854 | 3 | 220 | 91.6515 | 3 | 226.6667 | 70.2377 | 3 | 200 | 52.915 | 3 |
| 6 | 9.21E+08 | 7.1E+08 | 3 | 440 | 120 | 3 | 100 | 34.641 | 3 | 93.33333 | 50.3322 | 3 | 80 | 91.6515 | 3 | 26.66667 | 30.5505 | 3 |
| 8 | 1.85E+09 | 1.6E+08 | 3 | 280 | 105.83 | 3 | 40 | 34.641 | 3 | 6.666667 | 11.547 | 3 | 20 | 34.641 | 3 | 20 | 20 | 3 |
| 24 | 2.76E+09 | 7.2E+08 | 3 | 586.6667 | 231.805 | 3 | 0 | 0 | 3 | 0 | 0 | 3 | 0 | 0 | 3 | 0 | 0 | 3 |
| *E. coli* 25922 Site D | | | | | | | | | | | | | | | | | | |
| Timepoint | GC | | | x1 | | | x2 | | | x4 | | | x8 | | | x16 | | |
|  | Mean | SD | N | Mean | SD | N | Mean | SD | N | Mean | SD | N | Mean | SD | N | Mean | SD | N |
| 0 | 1566667 | 321455 | 3 | 2100000 | 556776 | 3 | 2133333.3 | 550757 | 3 | 1800000 | 300000 | 3 | 1866667 | 450925 | 3 | 1833333 | 115470 | 3 |
| 2 | 33000000 | 2645751 | 3 | 144000 | 79674.3 | 3 | 27333.333 | 6429.1 | 3 | 13333.33 | 3214.55 | 3 | 6033.333 | 1619.67 | 3 | 3833.333 | 208.167 | 3 |
| 4 | 82333333 | 2081666 | 3 | 24666.67 | 15534.9 | 3 | 2500 | 655.744 | 3 | 2400 | 888.819 | 3 | 1400 | 100 | 3 | 2366.667 | 208.167 | 3 |
| 6 | 2.57E+08 | 5773503 | 3 | 49666.67 | 42194 | 3 | 1633.3333 | 404.145 | 3 | 1900 | 458.258 | 3 | 1260 | 385.746 | 3 | 352 | 314.757 | 3 |
| 8 | 3.3E+08 | 2E+07 | 3 | 60666.67 | 22546.2 | 3 | 793.33333 | 272.274 | 3 | 953.3333 | 438.786 | 3 | 144 | 221.723 | 3 | 4.666667 | 1.1547 | 3 |
| 24 | 4E+08 | 0 | 3 | 2.27E+08 | 4.7E+07 | 3 | 1333.6667 | 1527.09 | 3 | 1 | 0 | 3 | 1 | 0 | 3 | 1 | 0 | 3 |
| *E. coli* 25922 Site E | | | | | | | | | | | | | | | | | | |
| Timepoint | GC | | | x1 | | | x2 | | | x4 | | | x8 | | | x16 | | |
|  | Mean | SD | N | Mean | SD | N | Mean | SD | N | Mean | SD | N | Mean | SD | N | Mean | SD | N |
| 0 | 13800000 | 1.1E+07 | 3 | 15313333 | 2.3E+07 | 3 | 24286667 | 2E+07 | 3 | 28666667 | 4725816 | 3 | 21200000 | 1.6E+07 | 3 | 29666667 | 2.4E+07 | 3 |
| 2 | 10733333 | 4818022 | 3 | 8233333 | 4750088 | 3 | 4533333.3 | 3061590 | 3 | 1866667 | 2294196 | 3 | 1606667 | 2420771 | 3 | 740000 | 1178474 | 3 |
| 4 | 2.02E+08 | 2.9E+08 | 3 | 6666667 | 1.2E+07 | 3 | 0 | 0 | 3 | 3333.333 | 5773.5 | 3 | 3333.333 | 5773.5 | 3 | 10000 | 10000 | 3 |
| 6 | 1.87E+09 | 3.2E+08 | 3 | 0 | 0 | 3 | 0 | 0 | 3 | 0 | 0 | 3 | 0 | 0 | 3 | 3333.333 | 5773.5 | 3 |
| 8 | 6.17E+09 | 8.5E+08 | 3 | 36666667 | 6.4E+07 | 3 | 0 | 0 | 3 | 0 | 0 | 3 | 0 | 0 | 3 | 0 | 0 | 3 |
| 24 | 6666.667 | 11547 | 3 | 1.15E+09 | 1.5E+09 | 3 | 0 | 0 | 3 | 0 | 0 | 3 | 0 | 0 | 3 | 0 | 0 | 3 |
| *E. coli* 25922 Site F | | | | | | | | | | | | | | | | | | |
| Timepoint | GC | | | x1 | | | x2 | | | x4 | | | x8 | | | x16 | | |
|  | Mean | SD | N | Mean | SD | N | Mean | SD | N | Mean | SD | N | Mean | SD | N | Mean | SD | N |
| 0 | 873333.3 | 76376.3 | 3 | 1356667 | 990168 | 3 | 896666.67 | 349905 | 3 | 800000 | 111355 | 3 | 1286667 | 795005 | 3 | 850000 | 70000 | 3 |
| 2 | 1846667 | 714516 | 3 | 990000 | 155885 | 3 | 1423333.3 | 845951 | 3 | 703333.3 | 185023 | 3 | 466666.7 | 97125.3 | 3 | 446666.7 | 105040 | 3 |
| 4 | 1.05E+08 | 8.3E+07 | 3 | 85333.33 | 25540.8 | 3 | 53000 | 8185.35 | 3 | 43000 | 14730.9 | 3 | 31333.33 | 8326.66 | 3 | 36666.67 | 9712.53 | 3 |
| 6 | 6.57E+08 | 4.7E+07 | 3 | 8800 | 2088.06 | 3 | 5100 | 1637.07 | 3 | 4933.333 | 1096.97 | 3 | 4400 | 1609.35 | 3 | 4133.333 | 1154.7 | 3 |
| 8 | 1.66E+09 | 5.9E+08 | 3 | 2733.333 | 650.641 | 3 | 1493.3333 | 666.433 | 3 | 1570 | 806.164 | 3 | 1136.667 | 553.203 | 3 | 1463.333 | 976.746 | 3 |
| 24 | 3.75E+09 | 2.5E+09 | 3 | 5.45E+08 | 7.7E+08 | 3 | 0 | 0 | 3 | 5 | 7.07107 | 3 | 0 | 0 | 3 | 0 | 0 | 3 |
| *E. coli* C1.55 Site A | | | | | | | | | | | | | | | | | | |
| Timepoint | GC | | | x1 | | | x2 | | | x4 | | | x8 | | | x16 | | |
|  | Mean | SD | N | Mean | SD | N | Mean | SD | N | Mean | SD | N | Mean | SD | N | Mean | SD | N |
| 0 | - | - | - | - | - | - | - | - | - | - | - | - | - | - | - | - | - | - |
| 2 | - | - | - | - | - | - | - | - | - | - | - | - | - | - | - | - | - | - |
| 4 | - | - | - | - | - | - | - | - | - | - | - | - | - | - | - | - | - | - |
| 6 | - | - | - | - | - | - | - | - | - | - | - | - | - | - | - | - | - | - |
| 8 | - | - | - | - | - | - | - | - | - | - | - | - | - | - | - | - | - | - |
| 24 | - | - | - | - | - | - | - | - | - | - | - | - | - | - | - | - | - | - |
| *E. coli* C1.55 Site B | | | | | | | | | | | | | | | | | | |
| Timepoint | GC |  |  | x1 |  |  | x2 |  |  | x4 |  |  | x8 |  |  | X16 |  |  |
|  | Mean | SD | N | Mean | SD | N | Mean | SD | N | Mean | SD | N | Mean | SD | N | Mean | SD | N |
| 0 | 800000 | 229129 | 3 | 683333 | 57735 | 3 | 783333 | 104083 | 3 | 750000 | 278388 | 3 | 850000 | 132288 | 3 | 733333 | 230940 | 3 |
| 2 | 15000000 | 2291288 | 3 | 96667 | 12583 | 3 | 11167 | 1893 | 3 | 7500 | 1323 | 3 | 5000 | 500 | 3 | 14667 | 17616 | 3 |
| 4 | 2.72E+08 | 3.1E+07 | 3 | 2933 | 1401 | 3 | 383 | 153 | 3 | 317 | 104 | 3 | 217 | 76 | 3 | 83 | 29 | 3 |
| 6 | 2.05E+09 | 2E+08 | 3 | 467 | 144 | 3 | 117 | 115 | 3 | 50.00333 | 50 | 3 | 200 | 264.571 | 3 | 17 | 28.8617 | 3 |
| 8 | 1.93E+09 | 3.8E+08 | 3 | 1050.003 | 1690.41 | 3 | 16.673333 | 28.8617 | 3 | 100.0033 | 132 | 3 | 1 | 0 | 3 | 33 | 58 | 3 |
| 24 | 8.83E+08 | 3.3E+08 | 3 | 1.5E+08 | 2.6E+08 | 3 | 1 | 0 | 3 | 17 | 28.8617 | 3 | 40067 | 69224 | 3 | 1 | 0 | 3 |
| *E. coli* C1.55 Site C | | | | | | | | | | | | | | | | | | |
| Timepoint | GC | | | x1 | | | x2 | | | x4 | | | x8 | | | x16 | | |
|  | Mean | SD | N | Mean | SD | N | Mean | SD | N | Mean | SD | N | Mean | SD | N | Mean | SD | N |
| 0 | 1226667 | 75718.8 | 3 | 1220000 | 34641 | 3 | 1293333.3 | 161658 | 3 | 1206667 | 117189 | 3 | 1233333 | 90185 | 3 | 441173.8 | 687511 | 3 |
| 2 | 17866667 | 1171893 | 3 | 378666.7 | 185487 | 3 | 10066.667 | 4163.33 | 3 | 4066.667 | 945.163 | 3 | 4000 | 346.41 | 3 | 1449.803 | 2215.2 | 3 |
| 4 | 4.4E+08 | 3.5E+07 | 3 | 7866.667 | 2571.64 | 3 | 886.66667 | 41.6333 | 3 | 346.6667 | 115.47 | 3 | 473.3333 | 70.2377 | 3 | 182.1903 | 254.369 | 3 |
| 6 | 1.28E+09 | 1.4E+08 | 3 | 593.3333 | 181.475 | 3 | 120 | 91.6515 | 3 | 86.66667 | 64.291 | 3 | 93.33333 | 11.547 | 3 | 35.96011 | 49.8701 | 3 |
| 8 | 1.35E+09 | 3.1E+08 | 3 | 160 | 52.915 | 3 | 53.333333 | 41.6333 | 3 | 33.33333 | 23.094 | 3 | 33.33333 | 23.094 | 3 | 19.80911 | 15.4312 | 3 |
| 24 | 1.68E+09 | 7.1E+08 | 3 | 87446.67 | 151168 | 3 | 0.6666667 | 0.57735 | 3 | 0.666667 | 0.57735 | 3 | 7 | 11.2694 | 3 | 7.089809 | 4.13545 | 3 |
| *E. coli* C1.55 Site D | | | | | | | | | | | | | | | | | | |
| Timepoint | GC | | | x1 | | | x2 | | | x4 | | | x8 | | | x16 | | |
|  | Mean | SD | N | Mean | SD | N | Mean | SD | N | Mean | SD | N | Mean | SD | N | Mean | SD | N |
| 0 | 1833333 | 305505 | 3 | 1566667 | 416333 | 3 | 1533333.3 | 321455 | 3 | 1766667 | 152753 | 3 | 1833333 | 152753 | 3 | 1733333 | 450925 | 3 |
| 2 | 18000000 | 2645751 | 3 | 310000 | 30000 | 3 | 71333.333 | 4509.25 | 3 | 24333.33 | 4725.82 | 3 | 27333.33 | 3055.05 | 3 | 24000 | 4358.9 | 3 |
| 4 | 97666667 | 1.1E+07 | 3 | 21466.67 | 16632.9 | 3 | 1023.3333 | 132.791 | 3 | 666.6667 | 46.188 | 3 | 586.6667 | 223.01 | 3 | 303.3333 | 170.098 | 3 |
| 6 | 2.6E+08 | 2E+07 | 3 | 1566.667 | 305.505 | 3 | 1733.3333 | 152.753 | 3 | 540 | 104.403 | 3 | 211.3333 | 336.668 | 3 | 23 | 16.6433 | 3 |
| 8 | 3.97E+08 | 5773503 | 3 | 7166.667 | 1738.77 | 3 | 5066.6667 | 472.582 | 3 | 169.6667 | 286.088 | 3 | 14.66667 | 8.5049 | 3 | 11.33333 | 3.21455 | 3 |
| 24 | 4E+08 | 0 | 3 | 2.5E+08 | 1.7E+07 | 3 | 15000000 | 4358899 | 3 | 6.333333 | 9.2376 | 3 | 1.666667 | 1.1547 | 3 | 5.333333 | 3.78594 | 3 |
| *E. col*i 25922 Site E | | | | | | | | | | | | | | | | | | |
| Timepoint | GC | | | x1 | | | x2 | | | x4 | | | x8 | | | x16 | | |
|  | Mean | SD | N | Mean | SD | N | Mean | SD | N | Mean | SD | N | Mean | SD | N | Mean | SD | N |
| 0 | 26333333 | 4041452 | 3 | 16000000 | 1.1E+07 | 3 | 19833333 | 1.5E+07 | 3 | 10433334 | 1.8E+07 | 3 | 28333333 | 3785939 | 3 | 19333335 | 1.7E+07 | 3 |
| 2 | 41300000 | 6E+07 | 3 | 5166667 | 1484363 | 3 | 2133333.3 | 1069268 | 3 | 4553333 | 7315465 | 3 | 253333.3 | 295014 | 3 | 206666.7 | 210792 | 3 |
| 4 | 1.36E+09 | 1.5E+09 | 3 | 166666.7 | 208167 | 3 | 0 | 0 | 3 | 13333.33 | 23094 | 3 | 3333.333 | 5773.5 | 3 | 6666.667 | 5773.5 | 3 |
| 6 | 2.6E+09 | 7.2E+08 | 3 | 100000 | 173205 | 3 | 0 | 0 | 3 | 3333.333 | 5773.5 | 3 | 0 | 0 | 3 | 0 | 0 | 3 |
| 8 | 6.53E+09 | 2.1E+09 | 3 | 366666.7 | 635085 | 3 | 0 | 0 | 3 | 0 | 0 | 3 | 0 | 0 | 3 | 0 | 0 | 3 |
| 24 | 240000 | 256320 | 3 | 9E+08 | 7.8E+08 | 3 | 0 | 0 | 3 | 0 | 0 | 3 | 0 | 0 | 3 | 0 | 0 | 3 |
| *E. coli* C1.55 Site F | | | | | | | | | | | | | | | | | | |
| Timepoint | GC | | | x1 | | | x2 | | | x4 | | | x8 | | | x16 | | |
|  | Mean | SD | N | Mean | SD | N | Mean | SD | N | Mean | SD | N | Mean | SD | N | Mean | SD | N |
| 0 | 1213333 | 342101 | 3 | 1130000 | 36055.5 | 3 | 1016666.7 | 251064 | 3 | 1420000 | 425676 | 3 | 7706667 | 1.1E+07 | 3 | 1196667 | 441626 | 3 |
| 2 | 2566667 | 680686 | 3 | 1660000 | 902663 | 3 | 996666.67 | 800021 | 3 | 1333333 | 755932 | 3 | 750000 | 206640 | 3 | 693333.3 | 260832 | 3 |
| 4 | 59666667 | 1.5E+07 | 3 | 253000 | 195952 | 3 | 149666.67 | 165385 | 3 | 98333.33 | 105368 | 3 | 61333.33 | 68156.7 | 3 | 105000 | 109659 | 3 |
| 6 | 5.03E+08 | 5.7E+07 | 3 | 97333.33 | 99042.1 | 3 | 30666.667 | 22854.6 | 3 | 18206.67 | 25817.1 | 3 | 22433.33 | 31672.4 | 3 | 14033.33 | 18159.4 | 3 |
| 8 | 1.64E+09 | 5.3E+08 | 3 | 32666.67 | 40501 | 3 | 14500 | 20352.1 | 3 | 6600 | 8140.64 | 3 | 10933.33 | 16542.5 | 3 | 13173.33 | 20647.8 | 3 |
| 24 | 2.33E+09 | 4E+08 | 3 | 3421000 | 5698614 | 3 | 10 | 17.3205 | 3 | 10 | 10 | 3 | 6.666667 | 11.547 | 3 | 10 | 10 | 3 |
| *E. coli* C1.62 Site A | | | | | | | | | | | | | | | | | | |
| Timepoint | GC | | | x1 | | | x2 | | | x4 | | | x8 | | | x16 | | |
|  | Mean | SD | N | Mean | SD | N | Mean | SD | N | Mean | SD | N | Mean | SD | N | Mean | SD | N |
| 0 | 600000 | 120000 | 3 | 546666.7 | 230940 | 3 | 640000 | 0 | 3 | 893333.3 | 161658 | 3 | 600000 | 105830 | 3 | 706666.7 | 23094 | 3 |
| 2 | 2818667 | 2400884 | 3 | 533866.7 | 923299 | 3 | 2400 | 692.82 | 3 | 266.6667 | 230.94 | 3 | 400 | 0 | 3 | 0 | 0 | 3 |
| 4 | 1.07E+08 | 1.3E+07 | 3 | 133.3333 | 230.94 | 3 | 0 | 0 | 3 | 0 | 0 | 3 | 0 | 0 | 3 | 0 | 0 | 3 |
| 6 | 2.27E+09 | 1E+09 | 3 | 266.6667 | 461.88 | 3 | 0 | 0 | 3 | 0 | 0 | 3 | 0 | 0 | 3 | 0 | 0 | 3 |
| 8 | - | - | - | - | - | - | - | - | - | - | - | - | - | - | - | - | - | - |
| 24 | 2.93E+09 | 3.7E+09 | 3 | 4.93E+09 | 1.8E+09 | 3 | 3.867E+09 | 8.3E+08 | 3 | 0 | 0 | 3 | 0 | 0 | 3 | 0 | 0 | 3 |
| *E. coli* C1.62 Site B | | | | | | | | | | | | | | | | | | |
| Timepoint | GC |  |  | x1 |  |  | x2 |  |  | x4 |  |  | x8 |  |  | X16 |  |  |
|  | Mean | SD | N | Mean | SD | N | Mean | SD | N | Mean | SD | N | Mean | SD | N | Mean | SD | N |
| 0 | 7.33E+05 | 104083 | 3 | 1.07E+06 | 104083 | 3 | 1.02E+06 | 301386 | 3 | 9.17E+05 | 236291 | 3 | 1.08E+06 | 57735 | 3 | 9.83E+05 | 57735 | 3 |
| 2 | 1.47E+07 | 1040833 | 3 | 4.00E+03 | 2646 | 3 | 6.67E+03 | 7217 | 3 | 3.33E+02 | 104 | 3 | 1.33E+02 | 58 | 3 | 0.00E+00 | 0 | 3 |
| 4 | 3.40E+08 | 9.7E+07 | 3 | 2.67E+02 | 126 | 3 | 6.70E+01 | 58 | 3 | 6.70E+01 | 58 | 3 | 8.30E+01 | 29 | 3 | 1.67E+02 | 289 | 3 |
| 6 | 1.65E+09 | 2.6E+08 | 3 | 3.00E+02 | 346 | 3 | 3.30E+01 | 58 | 3 | 3.67E+02 | 551 | 3 | 1.70E+01 | 28.8617 | 3 | 6.33E+02 | 1097 | 3 |
| 8 | 1.80E+09 | 2.6E+08 | 3 | 1.67E+02 | 175.589 | 3 | 1.67E+01 | 28.8617 | 3 | 1.00E+00 | 1 | 3 | 1.00E+00 | 1 | 3 | 1.00E+00 | 1 | 3 |
| 24 | 7.50E+08 | 1.3E+08 | 3 | 1.83E+08 | 1.6E+08 | 3 | 1.00E+00 | 1 | 3 | 1.00E+00 | 1 | 3 | 1.67E+04 | 28868 | 3 | 1.00E+00 | 1 | 3 |
| *E. coli* C1.62 Site C | | | | | | | | | | | | | | | | | | |
| Timepoint | GC | | | x1 | | | x2 | | | x4 | | | x8 | | | x16 | | |
|  | Mean | SD | N | Mean | SD | N | Mean | SD | N | Mean | SD | N | Mean | SD | N | Mean | SD | N |
| 0 | 626666.7 | 223010 | 3 | 540000 | 52915 | 3 | 546666.67 | 113725 | 3 | 553333.3 | 117189 | 3 | 613333.3 | 61101 | 3 | 700000 | 121655 | 3 |
| 2 | 8400000 | 2690725 | 3 | 9000 | 4275.51 | 3 | 8133.3333 | 3202.08 | 3 | 6000 | 1777.64 | 3 | 2740 | 3006.86 | 3 | 2113.333 | 1648.8 | 3 |
| 4 | 2.08E+08 | 4.4E+07 | 3 | 1386.667 | 725.075 | 3 | 1020 | 669.029 | 3 | 593.3333 | 375.411 | 3 | 220 | 140 | 3 | 120 | 120 | 3 |
| 6 | 1.36E+09 | 3.3E+08 | 3 | 13240 | 10260.1 | 3 | 340 | 196.977 | 3 | 226.6667 | 184.752 | 3 | 86.66667 | 83.2666 | 3 | 60 | 52.915 | 3 |
| 8 | 1.63E+09 | 2.6E+08 | 3 | 242266.7 | 247353 | 3 | 153.33333 | 98.6577 | 3 | 86.66667 | 30.5505 | 3 | 13.33333 | 23.094 | 3 | 20 | 20 | 3 |
| 24 | 2.4E+09 | 3.7E+08 | 3 | 2.13E+09 | 1E+09 | 3 | 1 | 0 | 3 | 1 | 0 | 3 | 1 | 0 | 3 | 7.333333 | 10.9697 | 3 |
| *E. coli* C1.62 Site D | | | | | | | | | | | | | | | | | | |
| Timepoint | GC | | | x1 | | | x2 | | | x4 | | | x8 | | | x16 | | |
|  | Mean | SD | N | Mean | SD | N | Mean | SD | N | Mean | SD | N | Mean | SD | N | Mean | SD | N |
| 0 | 1666667 | 378594 | 3 | 1633333 | 450925 | 3 | 1766666.7 | 251661 | 3 | 1600000 | 435890 | 3 | 1766667 | 378594 | 3 | 1533333 | 305505 | 3 |
| 2 | 19000000 | 1732051 | 3 | 13933.33 | 3579.57 | 3 | 7633.3333 | 1171.89 | 3 | 1866.667 | 472.582 | 3 | 533.3333 | 94.5163 | 3 | 7.666667 | 3.05505 | 3 |
| 4 | 1.09E+08 | 1.8E+07 | 3 | 940 | 103.923 | 3 | 553.33333 | 75.7188 | 3 | 14 | 2.64575 | 3 | 12.33333 | 9.29157 | 3 | 3.666667 | 0.57735 | 3 |
| 6 | 1.83E+08 | 2.5E+07 | 3 | 3700 | 1652.27 | 3 | 446.66667 | 28.8675 | 3 | 4.666667 | 1.52753 | 3 | 3.333333 | 0.57735 | 3 | 2 | 1 | 3 |
| 8 | 3.8E+08 | 3.5E+07 | 3 | 6633.333 | 416.333 | 3 | 2066.6667 | 680.686 | 3 | 2.666667 | 1.52753 | 3 | 1 | 0 | 3 | 1.333333 | 0.57735 | 3 |
| 24 | 4E+08 | 0 | 3 | 2E+08 | 7.5E+07 | 3 | 8000000 | 4331282 | 3 | 242 | 413.961 | 3 | 141.3333 | 241.334 | 3 | 1 | 0 | 3 |
| *E. coli* C1.62 Site E | | | | | | | | | | | | | | | | | | |
| Timepoint | GC | | | x1 | | | x2 | | | x4 | | | x8 | | | x16 | | |
|  | Mean | SD | N | Mean | SD | N | Mean | SD | N | Mean | SD | N | Mean | SD | N | Mean | SD | N |
| 0 | 545000 | 148492 | 2 | 610000 | 247588 | 3 | 666666.67 | 210792 | 3 | 640000 | 314325 | 3 | 600000 | 325115 | 3 | 800000 | 173494 | 3 |
| 2 | 13750000 | 7424621 | 2 | 2366667 | 4099187 | 3 | 0 | 0 | 3 | 0 | 0 | 3 | 0 | 0 | 3 | 0 | 0 | 3 |
| 4 | 6.26E+08 | 8.1E+08 | 2 | 8666667 | 1.5E+07 | 3 | 0 | 0 | 3 | 0 | 0 | 3 | 0 | 0 | 3 | 0 | 0 | 3 |
| 6 | 1.73E+09 | 2.4E+09 | 2 | 4.67E+08 | 8.1E+08 | 3 | 6666.6667 | 5773.5 | 3 | 0 | 0 | 3 | 0 | 0 | 3 | 0 | 0 | 3 |
| 8 | - | - | - | - | - | - | - | - | - | - | - | - | - | - | - | - | - | - |
| 24 | 3500000 | 2121320 | 2 | 1.41E+09 | 1.6E+09 | 3 | 1.58E+09 | 1.6E+09 | 3 | 0 | 0 | 3 | 0 | 0 | 3 | 0 | 0 | 3 |
| *E. coli* C1.62 Site F | | | | | | | | | | | | | | | | | | |
| Timepoint | GC | | | x1 | | | x2 | | | x4 | | | x8 | | | x16 | | |
|  | Mean | SD | N | Mean | SD | N | Mean | SD | N | Mean | SD | N | Mean | SD | N | Mean | SD | N |
| 0 | 863333.3 | 66583.3 | 3 | 803333.3 | 40414.5 | 3 | 893333.33 | 85049 | 3 | 806666.7 | 61101 | 3 | 726666.7 | 50332.2 | 3 | 660000 | 40000 | 3 |
| 2 | 920000 | 177764 | 3 | 260000 | 52915 | 3 | 176000 | 66090.8 | 3 | 76666.67 | 34151.6 | 3 | 136666.7 | 133095 | 3 | 90666.67 | 52548.4 | 3 |
| 4 | 61333333 | 3055050 | 3 | 73000 | 25865 | 3 | 42666.667 | 19008.8 | 3 | 32666.67 | 19502.1 | 3 | 41033.33 | 26905.5 | 3 | 51366.67 | 46184.4 | 3 |
| 6 | 7.9E+08 | 3.3E+08 | 3 | 594666.7 | 399231 | 3 | 16633.333 | 12023.4 | 3 | 15966.67 | 13229.6 | 3 | 11566.67 | 6819.34 | 3 | 12366.67 | 10209 | 3 |
| 8 | 1.86E+09 | 1.7E+09 | 3 | 1000000 | 0 | 3 | 18300 | 15330.7 | 3 | 6133.333 | 1625.83 | 3 | 4466.667 | 1504.44 | 3 | 4566.667 | 1703.92 | 3 |
| 24 | 1.86E+09 | 6E+08 | 3 | 1.12E+09 | 2.4E+08 | 3 | 1.38E+09 | 6.3E+08 | 3 | 5.7E+08 | 5E+08 | 3 | 6750 | 10614.7 | 3 | 170 | 115.326 | 3 |

Table S3- Different day source data all strains

| Different day replications | | | | | | | | | | | | | | | | | | |
| --- | --- | --- | --- | --- | --- | --- | --- | --- | --- | --- | --- | --- | --- | --- | --- | --- | --- | --- |
| *E. coli* 25922 Site A | | | | | | | | | | | | | | | | | | |
| Timepoint | GC | | | x1 | | | x2 | | | x4 | | | x8 | | | x16 | | |
|  | Mean | SD | N | Mean | SD | N | Mean | SD | N | Mean | SD | N | Mean | SD | N | Mean | SD | N |
| 0 | 667000 | 202073 | 3 | 3070 | 462 | 3 | 3600 | 0 | 3 | 3470 | 611 | 3 | 3870 | 231 | 3 | 3200 | 400 | 3 |
| 2 | 7830000 | 1892969 | 3 | 4400 | 693 | 3 | 3470 | 611 | 3 | 2270 | 231 | 3 | 0 | 0 | 3 | 0 | 0 | 3 |
| 4 | 1.73E+08 | 7.1E+07 | 3 | 32000 | 4000 | 3 | 267 | 231 | 3 | 0 | 0 | 3 | 0 | 0 | 3 | 0 | 0 | 3 |
| 6 | 6.17E+08 | 3.5E+08 | 3 | 1730000 | 1E+06 | 3 | 0 | 0 | 3 | 0 | 0 | 3 | 0 | 0 | 3 | 0 | 0 | 3 |
| 8 | 9.17E+08 | 2.9E+08 | 3 | 3.5E+07 | 8E+06 | 3 | 0 | 0 | 3 | 0 | 0 | 3 | 0 | 0 | 3 | 0 | 0 | 3 |
| 24 | 1.1E+09 | 4.3E+08 | 3 | 3.6E+07 | 1E+07 | 3 | 0 | 0 | 3 | 0 | 0 | 3 | 0 | 0 | 3 | 0 | 0 | 3 |
| *E. coli* 25922 Site B | | | | | | | | | | | | | | | | | | |
| Timepoint | GC | | | x1 | | | x2 | | | x4 | | | x8 | | | x16 | | |
|  | Mean | SD | N | Mean | SD | N | Mean | SD | N | Mean | SD | N | Mean | SD | N | Mean | SD | N |
| 0 | 633333.3 | 332916 | 3 | 633000 | 332916 | 3 | 400000 | 86602.5 | 3 | 400000 | 86602.5 | 3 | 617000 | 28868 | 3 | 700000 | 377492 | 3 |
| 2 | 56666.67 | 20816.7 | 3 | 56700 | 20817 | 3 | 26500 | 37672.9 | 3 | 4830 | 1040.83 | 3 | 1600 | 360.56 | 3 | 3420 | 2919.05 | 3 |
| 4 | 1550 | 576.628 | 3 | 1550 | 576.63 | 3 | 800 | 427.2 | 3 | 450 | 312.25 | 3 | 417 | 275.38 | 3 | 133 | 57.735 | 3 |
| 6 | 366.6667 | 340.343 | 3 | 367 | 340.34 | 3 | 117 | 125.826 | 3 | 83.3 | 28.8675 | 3 | 66.7 | 115.46 | 3 | 16.7 | 28.8617 | 3 |
| 8 | 15066.67 | 25923.1 | 3 | 15100 | 25923 | 3 | 50 | 49.995 | 3 | 0.01 | 0 | 3 | 16.7 | 28.862 | 3 | 0.01 | 0 | 3 |
| 24 | 84300 | 143504 | 3 | 84300 | 143504 | 3 | 500 | 476.964 | 3 | 63000 | 105708 | 3 | 8500 | 7466.6 | 3 | 0.01 | 0 | 3 |
| *E. coli* 25922 Site C | | | | | | | | | | | | | | | | | | |
| Timepoint | GC | | | x1 | | | x2 | | | x4 | | | x8 | | | x16 | | |
|  | Mean | SD | N | Mean | SD | N | Mean | SD | N | Mean | SD | N | Mean | SD | N | Mean | SD | N |
| 0 | 853333.3 | 211975 | 3 | 1180000 | 347000 | 3 | 1330000 | 469000 | 3 | 1240000 | 150997 | 3 | 1300000 | 231000 | 3 | 1473333 | 283784 | 3 |
| 2 | 1180000 | 346987 | 3 | 8210000 | 1E+07 | 3 | 31700 | 33300 | 3 | 7133.333 | 3590.73 | 3 | 5730 | 1700 | 3 | 5800 | 916.515 | 3 |
| 4 | 1333333 | 469184 | 3 | 12600 | 8340 | 3 | 2440 | 2910 | 3 | 760 | 623.859 | 3 | 520 | 347 | 3 | 386.6667 | 83.2666 | 3 |
| 6 | 1240000 | 150997 | 3 | 8030 | 9090 | 3 | 327 | 306 | 3 | 160 | 121.655 | 3 | 113 | 61.1 | 3 | 46.66667 | 30.5505 | 3 |
| 8 | 1300000 | 230651 | 3 | 120000 | 208000 | 3 | 73.3 | 75.7 | 3 | 86.66667 | 23.094 | 3 | 60 | 34.6 | 3 | 6.666667 | 11.547 | 3 |
| 24 | 1473333 | 283784 | 3 | 9.3E+08 | 2E+09 | 3 | 0 | 0 | 3 | 0 | 0 | 3 | 6.67 | 11.5 | 3 | 0 | 0 | 3 |
| *E. coli* 25922 Site D | | | | | | | | | | | | | | | | | | |
| Timepoint | GC | | | x1 | | | x2 | | | x4 | | | x8 | | | x16 | | |
|  | Mean | SD | N | Mean | SD | N | Mean | SD | N | Mean | SD | N | Mean | SD | N | Mean | SD | N |
| 0 | 1430000 | 252000 | 3 | 1760000 | 635000 | 3 | 1840000 | 559000 | 3 | 1430000 | 115000 | 3 | 1170000 | 208000 | 3 | 1330000 | 321455 | 3 |
| 2 | 17900000 | 1.6E+07 | 3 | 135000 | 71200 | 3 | 143000 | 224000 | 3 | 27000 | 21800 | 3 | 3600 | 2040 | 3 | 3770 | 611.01 | 3 |
| 4 | 38300000 | 4E+07 | 3 | 19400 | 13200 | 3 | 3180 | 1080 | 3 | 1830 | 513 | 3 | 1900 | 700 | 3 | 2000 | 435.89 | 3 |
| 6 | 2.53E+08 | 1.3E+08 | 3 | 41600 | 33000 | 3 | 1580 | 300 | 3 | 1330 | 208 | 3 | 1230 | 115 | 3 | 205 | 169.013 | 3 |
| 8 | 3.17E+08 | 3.2E+07 | 3 | 55800 | 17500 | 3 | 664 | 264 | 3 | 567 | 122 | 3 | 138 | 227 | 3 | 3.33 | 1.1547 | 3 |
| 24 | 4E+08 | 0 | 3 | 2.6E+08 | 6E+07 | 3 | 1840 | 1330 | 3 | 2 | 1.73 | 3 | 1.33 | 0.577 | 3 | 1 | 0 | 3 |
| *E. col*i 25922 Site E | | | | | | | | | | | | | | | | | | |
| Timepoint | GC | | | x1 | | | x2 | | | x4 | | | x8 | | | x16 | | |
|  | Mean | SD | N | Mean | SD | N | Mean | SD | N | Mean | SD | N | Mean | SD | N | Mean | SD | N |
| 0 | 1000 | 0 | 2 | 1570000 | 1E+06 | 3 | 1710000 | 1466470 | 3 | 2640000 | 121655 | 3 | 3200000 | 484974 | 3 | 10500000 | 1281718 | 3 |
| 2 | 1000 | 0 | 2 | 1.2E+07 | 3E+06 | 3 | 6470000 | 2655811 | 3 | 3270000 | 2136196 | 3 | 2940000 | 2528794 | 3 | 4220000 | 1200889 | 3 |
| 4 | 10000 | 0 | 2 | 2670000 | 5E+06 | 3 | 1500000 | 2598076 | 3 | 3330 | 5773.5 | 3 | 3330 | 5773.5 | 3 | 10000 | 5773.5 | 3 |
| 6 | 1000000 | 0 | 2 | 1500000 | 3E+06 | 3 | 0 | 0 | 3 | 0 | 0 | 3 | 3330 | 5773.5 | 3 | 10000 | 5773.5 | 3 |
| 8 | 1000000 | 0 | 2 | 5270000 | 6E+06 | 3 | 0 | 0 | 3 | 0 | 0 | 3 | 0 | 0 | 3 | 0 | 0 | 3 |
| 24 | 10000000 | 0 | 2 | 2.2E+08 | 4E+08 | 3 | 0 | 0 | 3 | 0 | 0 | 3 | 0 | 0 | 3 | 0 | 0 | 3 |
| *E. coli* 25922 Site F | | | | | | | | | | | | | | | | | | |
| Timepoint | GC | | | x1 | | | x2 | | | x4 | | | x8 | | | x16 | | |
|  | Mean | SD | N | Mean | SD | N | Mean | SD | N | Mean | SD | N | Mean | SD | N | Mean | SD | N |
| 0 | 873000 | 76400 | 3 | 1360000 | 990000 | 3 | 1710000 | 1466470 | 3 | 800000 | 111000 | 3 | 1290000 | 795000 | 3 | 850000 | 70000 | 3 |
| 2 | 1850000 | 715000 | 3 | 990000 | 156000 | 3 | 6470000 | 2655811 | 3 | 703000 | 185000 | 3 | 467000 | 97100 | 3 | 447000 | 105040 | 3 |
| 4 | 1.05E+08 | 8.3E+07 | 3 | 85300 | 25500 | 3 | 1500000 | 2598076 | 3 | 43000 | 14700 | 3 | 98000 | 107000 | 3 | 36700 | 9712.54 | 3 |
| 6 | 6.57E+08 | 4.7E+07 | 3 | 8800 | 2090 | 3 | 0 | 0 | 3 | 4930 | 1100 | 3 | 4400 | 1610 | 3 | 3930 | 1512.66 | 3 |
| 8 | 1.66E+09 | 5.9E+08 | 3 | 9400 | 11500 | 3 | 0 | 0 | 3 | 1560 | 807 | 3 | 1140 | 553 | 3 | 1460 | 976.746 | 3 |
| 24 | 4.2E+09 | 1.9E+09 | 3 | 3.7E+08 | 6E+08 | 3 | 0 | 0 | 3 | 4 | 5.2 | 3 | 1 | 0 | 3 | 13700000 | 2.4E+07 | 3 |
| *E. coli* C1.55 Site A | | | | | | | | | | | | | | | | | | |
| Timepoint | GC | | | x1 | | | x2 | | | x4 | | | x8 | | | x16 | | |
|  | Mean | SD | N | Mean | SD | N | Mean | SD | N | Mean | SD | N | Mean | SD | N | Mean | SD | N |
| 0 | - | - | - | - | - | - | - | - | - | - | - | - | - | - | - | - | - | - |
| 2 | - | - | - | - | - | - | - | - | - | - | - | - | - | - | - | - | - | - |
| 4 | - | - | - | - | - | - | - | - | - | - | - | - | - | - | - | - | - | - |
| 6 | - | - | - | - | - | - | - | - | - | - | - | - | - | - | - | - | - | - |
| 8 | - | - | - | - | - | - | - | - | - | - | - | - | - | - | - | - | - | - |
| 24 | - | - | - | - | - | - | - | - | - | - | - | - | - | - | - | - | - | - |
| *E. coli* C1.55 Site B | | | | | | | | | | | | | | | | | | |
| Timepoint | GC |  |  | x1 |  |  | x2 |  |  | x4 |  |  | x8 |  |  | X16 |  |  |
|  | Mean | SD | N | Mean | SD | N | Mean | SD | N | Mean | SD | N | Mean | SD | N | Mean | SD | N |
| 0 | 800000 | 229129 | 3 | 683333 | 57735 | 3 | 783333 | 104083 | 3 | 750000 | 278388 | 3 | 850000 | 132288 | 3 | 733333 | 230940 | 3 |
| 2 | 15000000 | 2291288 | 3 | 96667 | 12583 | 3 | 11167 | 1893 | 3 | 7500 | 1323 | 3 | 5000 | 500 | 3 | 14667 | 17616 | 3 |
| 4 | 2.72E+08 | 3.1E+07 | 3 | 2933 | 1401 | 3 | 383 | 153 | 3 | 317 | 104 | 3 | 217 | 76 | 3 | 83 | 29 | 3 |
| 6 | 2.05E+09 | 2E+08 | 3 | 467 | 144 | 3 | 117 | 115 | 3 | 50.00333 | 50 | 3 | 200 | 264.571 | 3 | 17 | 28.8617 | 3 |
| 8 | 1.93E+09 | 3.8E+08 | 3 | 1050.003 | 1690.41 | 3 | 16.673333 | 28.8617 | 3 | 100.0033 | 132 | 3 | 1 | 0 | 3 | 33 | 58 | 3 |
| 24 | 8.83E+08 | 3.3E+08 | 3 | 1.5E+08 | 2.6E+08 | 3 | 1 | 0 | 3 | 17 | 28.8617 | 3 | 40067 | 69224 | 3 | 1 | 0 | 3 |
| *E. coli* C1.55 Site C | | | | | | | | | | | | | | | | | | |
| Timepoint | GC | | | x1 | | | x2 | | | x4 | | | x8 | | | x16 | | |
|  | Mean | SD | N | Mean | SD | N | Mean | SD | N | Mean | SD | N | Mean | SD | N | Mean | SD | N |
| 0 | 1226667 | 75718.8 | 3 | 1220000 | 34641 | 3 | 1293333.3 | 161658 | 3 | 1206667 | 117189 | 3 | 1233333 | 90185 | 3 | 441173.8 | 687511 | 3 |
| 2 | 17866667 | 1171893 | 3 | 378666.7 | 185487 | 3 | 10066.667 | 4163.33 | 3 | 4066.667 | 945.163 | 3 | 4000 | 346.41 | 3 | 1449.803 | 2215.2 | 3 |
| 4 | 4.4E+08 | 3.5E+07 | 3 | 7866.667 | 2571.64 | 3 | 886.66667 | 41.6333 | 3 | 346.6667 | 115.47 | 3 | 473.3333 | 70.2377 | 3 | 182.1903 | 254.369 | 3 |
| 6 | 1.28E+09 | 1.4E+08 | 3 | 593.3333 | 181.475 | 3 | 120 | 91.6515 | 3 | 86.66667 | 64.291 | 3 | 93.33333 | 11.547 | 3 | 35.96011 | 49.8701 | 3 |
| 8 | 1.35E+09 | 3.1E+08 | 3 | 160 | 52.915 | 3 | 53.333333 | 41.6333 | 3 | 33.33333 | 23.094 | 3 | 33.33333 | 23.094 | 3 | 19.80911 | 15.4312 | 3 |
| 24 | 1.68E+09 | 7.1E+08 | 3 | 87446.67 | 151168 | 3 | 0.6666667 | 0.57735 | 3 | 0.666667 | 0.57735 | 3 | 7 | 11.2694 | 3 | 7.089809 | 4.13545 | 3 |
| *E. coli* C1.55 Site D | | | | | | | | | | | | | | | | | | |
| Timepoint | GC | | | x1 | | | x2 | | | x4 | | | x8 | | | x16 | | |
|  | Mean | SD | N | Mean | SD | N | Mean | SD | N | Mean | SD | N | Mean | SD | N | Mean | SD | N |
| 0 | 1833333 | 305505 | 3 | 1566667 | 416333 | 3 | 1533333.3 | 321455 | 3 | 1766667 | 152753 | 3 | 1833333 | 152753 | 3 | 1733333 | 450925 | 3 |
| 2 | 18000000 | 2645751 | 3 | 310000 | 30000 | 3 | 71333.333 | 4509.25 | 3 | 24333.33 | 4725.82 | 3 | 27333.33 | 3055.05 | 3 | 24000 | 4358.9 | 3 |
| 4 | 97666667 | 1.1E+07 | 3 | 21466.67 | 16632.9 | 3 | 1023.3333 | 132.791 | 3 | 666.6667 | 46.188 | 3 | 586.6667 | 223.01 | 3 | 303.3333 | 170.098 | 3 |
| 6 | 2.6E+08 | 2E+07 | 3 | 1566.667 | 305.505 | 3 | 1733.3333 | 152.753 | 3 | 540 | 104.403 | 3 | 211.3333 | 336.668 | 3 | 23 | 16.6433 | 3 |
| 8 | 3.97E+08 | 5773503 | 3 | 7166.667 | 1738.77 | 3 | 5066.6667 | 472.582 | 3 | 169.6667 | 286.088 | 3 | 14.66667 | 8.5049 | 3 | 11.33333 | 3.21455 | 3 |
| 24 | 4E+08 | 0 | 3 | 2.5E+08 | 1.7E+07 | 3 | 15000000 | 4358899 | 3 | 6.333333 | 9.2376 | 3 | 1.666667 | 1.1547 | 3 | 5.333333 | 3.78594 | 3 |
| *E. coli* 25922 Site E | | | | | | | | | | | | | | | | | | |
| Timepoint | GC | | | x1 | | | x2 | | | x4 | | | x8 | | | x16 | | |
|  | Mean | SD | N | Mean | SD | N | Mean | SD | N | Mean | SD | N | Mean | SD | N | Mean | SD | N |
| 0 | 26333333 | 4041452 | 3 | 16000000 | 1.1E+07 | 3 | 19833333 | 1.5E+07 | 3 | 10433334 | 1.8E+07 | 3 | 28333333 | 3785939 | 3 | 19333335 | 1.7E+07 | 3 |
| 2 | 41300000 | 6E+07 | 3 | 5166667 | 1484363 | 3 | 2133333.3 | 1069268 | 3 | 4553333 | 7315465 | 3 | 253333.3 | 295014 | 3 | 206666.7 | 210792 | 3 |
| 4 | 1.36E+09 | 1.5E+09 | 3 | 166666.7 | 208167 | 3 | 0 | 0 | 3 | 13333.33 | 23094 | 3 | 3333.333 | 5773.5 | 3 | 6666.667 | 5773.5 | 3 |
| 6 | 2.6E+09 | 7.2E+08 | 3 | 100000 | 173205 | 3 | 0 | 0 | 3 | 3333.333 | 5773.5 | 3 | 0 | 0 | 3 | 0 | 0 | 3 |
| 8 | 6.53E+09 | 2.1E+09 | 4 | 366666.7 | 635085 | 3 | 0 | 0 | 3 | 0 | 0 | 3 | 0 | 0 | 3 | 0 | 0 | 3 |
| 24 | 240000 | 256320 | 3 | 9E+08 | 7.8E+08 | 3 | 0 | 0 | 3 | 0 | 0 | 3 | 0 | 0 | 3 | 0 | 0 | 3 |
| *E. coli* C1.55 Site F | | | | | | | | | | | | | | | | | | |
| Timepoint | GC | | | x1 | | | x2 | | | x4 | | | x8 | | | x16 | | |
|  | Mean | SD | N | Mean | SD | N | Mean | SD | N | Mean | SD | N | Mean | SD | N | Mean | SD | N |
| 0 | 1213333 | 342101 | 3 | 1130000 | 36055.5 | 3 | 1016666.7 | 251064 | 3 | 1420000 | 425676 | 3 | 7706667 | 1.1E+07 | 3 | 1196667 | 441626 | 3 |
| 2 | 2566667 | 680686 | 3 | 1660000 | 902663 | 3 | 996666.67 | 800021 | 3 | 1333333 | 755932 | 3 | 750000 | 206640 | 3 | 693333.3 | 260832 | 3 |
| 4 | 59666667 | 1.5E+07 | 3 | 253000 | 195952 | 3 | 149666.67 | 165385 | 3 | 98333.33 | 105368 | 3 | 61333.33 | 68156.7 | 3 | 105000 | 109659 | 3 |
| 6 | 5.03E+08 | 5.7E+07 | 3 | 97333.33 | 99042.1 | 3 | 30666.667 | 22854.6 | 3 | 18206.67 | 25817.1 | 3 | 22433.33 | 31672.4 | 3 | 14033.33 | 18159.4 | 3 |
| 8 | 1.64E+09 | 5.3E+08 | 3 | 32666.67 | 40501 | 3 | 14500 | 20352.1 | 3 | 6600 | 8140.64 | 3 | 10933.33 | 16542.5 | 3 | 13173.33 | 20647.8 | 3 |
| 24 | 2.33E+09 | 4E+08 | 3 | 3421000 | 5698614 | 3 | 10 | 17.3205 | 3 | 10 | 10 | 3 | 6.666667 | 11.547 | 3 | 10 | 10 | 3 |
| *E. coli* C1.62 Site A | | | | | | | | | | | | | | | | | | |
| Timepoint | GC | | | x1 | | | x2 | | | x4 | | | x8 | | | x16 | | |
|  | Mean | SD | N | Mean | SD | N | Mean | SD | N | Mean | SD | N | Mean | SD | N | Mean | SD | N |
| 0 | 600000 | 120000 | 3 | 546666.7 | 230940 | 3 | 640000 | 0 | 3 | 893333.3 | 161658 | 3 | 600000 | 105830 | 3 | 706666.7 | 23094 | 3 |
| 2 | 2818667 | 2400884 | 3 | 533866.7 | 923299 | 3 | 2400 | 692.82 | 3 | 266.6667 | 230.94 | 3 | 400 | 0 | 3 | 0 | 0 | 3 |
| 4 | 1.07E+08 | 1.3E+07 | 3 | 133.3333 | 230.94 | 3 | 0 | 0 | 3 | 0 | 0 | 3 | 0 | 0 | 3 | 0 | 0 | 3 |
| 6 | 2.27E+09 | 1E+09 | 3 | 266.6667 | 461.88 | 3 | 0 | 0 | 3 | 0 | 0 | 3 | 0 | 0 | 3 | 0 | 0 | 3 |
| 8 | - | - | - | - | - | - | - | - | - | - | - | - | - | - | - | - | - | - |
| 24 | 2.93E+09 | 3.7E+09 | 3 | 4.93E+09 | 1.8E+09 | 3 | 3.867E+09 | 8.3E+08 | 3 | 0 | 0 | 3 | 0 | 0 | 3 | 0 | 0 | 3 |
| *E. coli* C1.62 Site B | | | | | | | | | | | | | | | | | | |
| Timepoint | GC |  |  | x1 |  |  | x2 |  |  | x4 |  |  | x8 |  |  | X16 |  |  |
|  | Mean | SD | N | Mean | SD | N | Mean | SD | N | Mean | SD | N | Mean | SD | N | Mean | SD | N |
| 0 | 6.50E+05 | 278388 | 3 | 7.67E+05 | 275379 | 3 | 9.17E+05 | 256580 | 3 | 4.83E+05 | 230940 | 3 | 5.17E+05 | 208167 | 3 | 8.83E+05 | 160728 | 3 |
| 2 | 7.50E+06 | 866025 | 3 | 1.83E+03 | 763.763 | 3 | 1.30E+03 | 200 | 3 | 3.67E+02 | 57.735 | 3 | 1.83E+02 | 275.375 | 3 | 1.00E-02 | 0 | 3 |
| 4 | 1.92E+08 | 7.5E+07 | 3 | 2.83E+02 | 115.47 | 3 | 8.33E+01 | 76.3708 | 3 | 5.00E+01 | 49.995 | 3 | 1.00E-02 | 0 | 3 | 1.00E-02 | 0 | 3 |
| 6 | 7.17E+08 | 1.3E+08 | 3 | 3.18E+03 | 5470.45 | 3 | 5.00E+01 | 49.995 | 3 | 1.00E-02 | 0 | 3 | 1.00E-02 | 0 | 3 | 1.00E-02 | 0 | 3 |
| 8 | 7.83E+08 | 7.6E+07 | 3 | 2.33E+05 | 404116 | 3 | 1.00E-02 | 0 | 3 | 1.00E-02 | 0 | 3 | 1.00E-02 | 0 | 3 | 1.00E-02 | 0 | 3 |
| 24 | 9.67E+08 | 4.3E+08 | 3 | 3.67E+08 | 4E+08 | 3 | 1.00E+05 | 173205 | 3 | 1.00E-02 | 0 | 3 | 1.00E-02 | 0 | 3 | 1.00E-02 | 0 | 3 |
| *E. coli* C1.62 Site C | | | | | | | | | | | | | | | | | | |
| Timepoint | GC | | | x1 | | | x2 | | | x4 | | | x8 | | | x16 | | |
|  | Mean | SD | N | Mean | SD | N | Mean | SD | N | Mean | SD | N | Mean | SD | N | Mean | SD | N |
| 0 | 900000 | 250599 | 3 | 1086667 | 102632 | 3 | 1000000 | 87178 | 3 | 906666.7 | 236925 | 3 | 940000 | 60000 | 3 | 1060000 | 124900 | 3 |
| 2 | 10004467 | 8710113 | 3 | 1840 | 366.606 | 3 | 1413.3333 | 460.579 | 3 | 613.3333 | 166.533 | 3 | 106.6667 | 11.547 | 3 | 20 | 20 | 3 |
| 4 | 3.53E+08 | 3.1E+07 | 3 | 1793.333 | 1050.78 | 3 | 613.33333 | 560.119 | 3 | 53.33333 | 23.094 | 3 | 6.666667 | 11.547 | 3 | 0 | 0 | 3 |
| 6 | 1.09E+09 | 8.4E+08 | 3 | 44000 | 42142.6 | 3 | 16940 | 16070.9 | 3 | 13.33333 | 11.547 | 3 | 6.666667 | 11.547 | 3 | 0 | 0 | 3 |
| 8 | 1.72E+09 | 1.2E+08 | 3 | 2790000 | 3235939 | 3 | 393333.33 | 447363 | 3 | 0 | 0 | 3 | 0 | 0 | 3 | 0 | 0 | 3 |
| 24 | 2.15E+09 | 4.2E+08 | 3 | 1.89E+09 | 6.3E+08 | 3 | 433333333 | 5.4E+08 | 3 | 0 | 0 | 3 | 0 | 0 | 3 | 0 | 0 | 3 |
| *E. coli* C1.62 Site D | | | | | | | | | | | | | | | | | | |
| Timepoint | GC | | | x1 | | | x2 | | | x4 | | | x8 | | | x16 | | |
|  | Mean | SD | N | Mean | SD | N | Mean | SD | N | Mean | SD | N | Mean | SD | N | Mean | SD | N |
| 0 | 1500000 | 360555 | 3 | 1633333 | 351188 | 3 | 1866666.7 | 305505 | 3 | 1800000 | 173205 | 3 | 1633333 | 230940 | 3 | 1400000 | 346410 | 3 |
| 2 | 15000000 | 3605551 | 3 | 13666.67 | 2516.61 | 3 | 8166.6667 | 115.47 | 3 | 1733.333 | 57.735 | 3 | 480 | 34.641 | 3 | 22.66667 | 24.0069 | 3 |
| 4 | 89333333 | 7767453 | 3 | 1256.667 | 653.937 | 3 | 566.66667 | 83.2666 | 3 | 21 | 18.1934 | 3 | 16 | 6.55744 | 3 | 6 | 3.60555 | 3 |
| 6 | 2.23E+08 | 9.3E+07 | 3 | 4133.333 | 1205.54 | 3 | 316.66667 | 162.891 | 3 | 8.333333 | 3.05505 | 3 | 2.666667 | 1.52753 | 3 | 4 | 1.73205 | 3 |
| 8 | 3.33E+08 | 1.2E+08 | 3 | 44833.33 | 34671.1 | 3 | 1566.6667 | 305.505 | 3 | 4.666667 | 3.05505 | 3 | 2.666667 | 2.88675 | 3 | 2 | 1 | 3 |
| 24 | 4E+08 | 0 | 3 | 2.57E+08 | 8.1E+07 | 3 | 60666667 | 7.7E+07 | 3 | 4.333333 | 0.57735 | 3 | 141 | 241.622 | 3 | 1 | 0 | 3 |
| *E. coli* C1.62 Site E | | | | | | | | | | | | | | | | | | |
| Timepoint | GC | | | x1 | | | x2 | | | x4 | | | x8 | | | x16 | | |
|  | Mean | SD | N | Mean | SD | N | Mean | SD | N | Mean | SD | N | Mean | SD | N | Mean | SD | N |
| 0 | 550000 | 132288 | 3 | 513333.3 | 80829 | 3 | 556666.67 | 20816.7 | 3 | 520000 | 111355 | 3 | 410000 | 55677.6 | 3 | 480000 | 252389 | 3 |
| 2 | 11600000 | 6618912 | 3 | 0 | 0 | 3 | 0 | 0 | 3 | 0 | 0 | 3 | 0 | 0 | 3 | 0 | 0 | 3 |
| 4 | 6.67E+08 | 5E+08 | 3 | 0 | 0 | 3 | 0 | 0 | 3 | 0 | 0 | 3 | 0 | 0 | 3 | 0 | 0 | 3 |
| 6 | 1.4E+09 | 1.8E+09 | 3 | 0 | 0 | 3 | 0 | 0 | 3 | 0 | 0 | 3 | 0 | 0 | 3 | 0 | 0 | 3 |
| 8 | - | - | - | - | - | - | - | - | - | - | - | - | - | - | - | - | - | - |
| 24 | 40000000 | 1.7E+07 | 3 | 2.08E+09 | 1.1E+09 | 3 | 2.063E+09 | 1.1E+09 | 3 | 0 | 0 | 3 | 0 | 0 | 3 | 0 | 0 | 3 |
| *E. coli* C1.62 Site F | | | | | | | | | | | | | | | | | | |
| Timepoint | GC | | | x1 | | | x2 | | | x4 | | | x8 | | | x16 | | |
|  | Mean | SD | N | Mean | SD | N | Mean | SD | N | Mean | SD | N | Mean | SD | N | Mean | SD | N |
| 0 | 876666.7 | 230940 | 3 | 910000 | 173494 | 3 | 883333.33 | 168622 | 3 | 870000 | 196723 | 3 | 1370000 | 810987 | 3 | 636666.7 | 556447 | 3 |
| 2 | 3006667 | 1890009 | 3 | 156333.3 | 166584 | 3 | 131066.67 | 145573 | 3 | 90333.33 | 98083.3 | 3 | 88733.33 | 84574.3 | 3 | 42133.33 | 39131.2 | 3 |
| 4 | 1.43E+08 | 1.3E+08 | 3 | 21233.33 | 17957.3 | 3 | 26933.333 | 22654.2 | 3 | 11043.33 | 15670.9 | 3 | 19383.33 | 16924.6 | 3 | 5676.667 | 4940 | 3 |
| 6 | 7.43E+08 | 2.7E+08 | 3 | 12433.33 | 12890.4 | 3 | 4660 | 4260.14 | 3 | 3736.667 | 3905.9 | 3 | 2570 | 2545.13 | 3 | 1213.333 | 1656.06 | 3 |
| 8 | 1.13E+09 | 1.1E+09 | 3 | 31733.33 | 30571.4 | 3 | 5700 | 2722.13 | 3 | 3436.667 | 3086.91 | 3 | 1446.667 | 2217.03 | 3 | 700 | 1126.54 | 3 |
| 24 | 1.54E+09 | 4.8E+08 | 3 | 1.37E+09 | 5.6E+08 | 3 | 643334600 | 5.9E+08 | 3 | 400 | 649.692 | 3 | 116.6667 | 160.728 | 3 | 36.66667 | 63.5085 | 3 |

Table S4 – Intra-site Friedman analysis

| Intra-site analysis | | | | | | | | |
| --- | --- | --- | --- | --- | --- | --- | --- | --- |
| SD rapid kill Friedman scores | | | |  | DD rapid kill Friedman scores | | | |
| Site | 25922 | C1.55 | C1.62 |  | Site | 25922 | C1.55 | C1.62 |
|  | ATCC | ESBL | OXA-48 |  |  | ATCC | ESBL | OXA-48 |
|  | E.coli | E.coli | E.coli |  |  | E.coli | E.coli | E.coli |
| A | 22.97 | 0 | 14.76 |  | A | 23.79 | 0 | 14.76 |
| B | 19.57 | 16.57 | 12.41 |  | B | 19.25 | 17 | 15 |
| C | 26.95 | 26.64 | 29.04 |  | C | 28.07 | 27.23 | 29.04 |
| D | 42.68 | 45.61 | 49.27 |  | D | 18.25 | 17.85 | 19.75 |
| E | 14.83 | 17.92 | 8.57 |  | E | 17.19 | 20.21 | 8.57 |
| F | 23.73 | 21.33 | 21.69 |  | F | 29.7 | 16.2 | 20.2 |
| SD all data points Friedman scores | | | |  | DD all data points Friedman scores | | | |
| Site | 25922 | C1.55 | C1.62 |  | Site | 25922 | C1.55 | C1.62 |
|  | ATCC | ESBL | OXA-48 |  |  | ATCC | ESBL | OXA-48 |
|  | E.coli | E.coli | E.coli |  |  | E.coli | E.coli | E.coli |
| A | 23.79 | 0 | 14.76 |  | A | 19.61 | 0 | 0 |
| B | 19.25 | 17 | 15 |  | B | 28.11 | 28.84 | 27.19 |
| C | 28.07 | 27.23 | 29.04 |  | C | 35.904 | 43.11 | 52.69 |
| D | 18.25 | 17.85 | 19.75 |  | D | 18.25 | 17.85 | 13.75 |
| E | 17.19 | 20.21 | 8.57 |  | E | 11.07 | 26.9 | 9.15 |
| F | 29.7 | 16.2 | 20.2 |  | F | 16.57 | 14.86 | 25.83 |

Table S5 - Inter-site Friedman analysis

| Inter-site analysis | | | | | |
| --- | --- | --- | --- | --- | --- |
| SD 25922 | | | | | |
|  | MIC multiple | | | | |
|  | x1 | x2 | x4 | x8 | x16 |
| Rapid kill | 6.211811 | 10.90953 | 11.49371 | 16.18337 | 12.08325 |
| All data points | 4.831004 | 12.64598 | 16.37971 | 19.76531 | 10.63174 |
| DD 25922 | | | | | |
|  | MIC multiple | | | | |
|  | x1 | x2 | x4 | x8 | x16 |
| Rapid kill | 5.662376 | 8.726772 | 11.76637 | 13.47619 | 12.71429 |
| All data points | 6.118264 | 10.87148 | 7.942323 | 10.45771 | 8.453903 |
| SD C1.55 | | | | | |
|  | MIC multiple | | | | |
|  | x1 | x2 | x4 | x8 | x16 |
| Rapid kill | 9.52 | 8.47619 | 13.66667 | 14.2381 | 11.57143 |
| All data points | 6.195129 | 9.578727 | 18 | 16.02381 | 17.35714 |
| DD C1.55 | | | | | |
|  | MIC multiple | | | | |
|  | x1 | x2 | x4 | x8 | x16 |
| Rapid kill | 12.714 | 8.47619 | 13.6666 | 14.238 | 13.476 |
| All data points | 8.963 | 15.61904 | 11.02369 | 15.3809 | 16.976 |
| SD C1.62 | | | | | |
|  | MIC multiple | | | | |
|  | x1 | x2 | x4 | x8 | x16 |
| Rapid kill | 5.095 | 8.85714 | 7.047618 | 10.0476 | 3.80952 |
| All data points | 6.45238 | 10.738 | 18.9285 | 21.61 | 14.73809 |
| DD C1.62 | | | | | |
|  | MIC multiple | | | | |
|  | x1 | x2 | x4 | x8 | x16 |
| Rapid kill | 10.80952 | 10.19048 | 9.142857 | 12.5714 | 9.0476 |
| All data points | 9.83048 | 12.59524 | 18.3809 | 22.38095 | 20.16666 |
| SD Site C and D only 25922 | | | | | |
|  | MIC multiple | | | | |
|  | x1 | x2 | x4 | x8 | x16 |
| Rapid kill | 10.33 | 10.71428 | 10.71429 | 10.33 | 10.333 |
| All data points | 6.54 | 9.61 | 12.62 | 15.56 | 17.3809 |
